# A flexible Bayesian approach to estimating size-structured matrix population models

**DOI:** 10.1101/2021.07.16.452528

**Authors:** Jann Paul Mattern, Kristof Glauninger, Gregory L. Britten, John Casey, Sangwon Hyun, Zhen Wu, E Virginia Armbrust, Zaid Harchaoui, François Ribalet

## Abstract

The rates of cell growth, division, and carbon loss of microbial populations are key parameters for understanding how organisms interact with their environment and how they contribute to the carbon cycle. However, the invasive nature of current analytical methods has hindered efforts to reliably quantify these parameters. In recent years, size-structured matrix population models (MPMs) have gained popularity for estimating rate parameters of microbial populations by mechanistically describing changes in microbial cell size distributions over time. And yet, the construction, analysis, and biological interpretation of these models are underdeveloped, as current implementations do not adequately constrain or assess the biological feasibility of parameter values, leading to inference which may provide a good fit to observed size distributions but does not necessarily reflect realistic physiological dynamics. Here we present a flexible Bayesian extension of size-structured MPMs for testing underlying assumptions describing the dynamics of a marine phytoplankton population over the day-night cycle. Our Bayesian framework takes prior scientific knowledge into account and generates biologically interpretable results. Using data from an exponentially growing laboratory culture of the cyanobacterium *Prochlorococcus*, we herein demonstrate the performance improvements of our approach over current models and isolate previously ignored biological processes, such as respiratory and exudative carbon losses, as critical parameters for the modeling of microbial population dynamics. The results demonstrate that this modeling framework can provide deeper insights into microbial population dynamics provided by flow-cytometry time-series data.

**Author summary:** Identifying the growth and population dynamics of marine microorganisms in their natural habitat is crucial to understanding the flow of carbon in the oceans but remains a grand challenge due to the invasive nature of current measurement methods. As time-series observations of population size structure have become more commonplace in aquatic environments, matrix population models (MPMs), which aim to mechanistically describe the change in size structure of these populations over time, have gained in popularity over the last decade. However, the underlying assumptions and behavior of MPMs have not been adequately scrutinized, and parameter values are difficult to interpret biologically, leading to inference that may not reflect plausible physiological dynamics. Here, we develop a Bayesian extension of the MPM framework to examine biological assumptions, improve interpretability of model output, and account for additional biological processes. We evaluated the performance of our models on a publicly available dataset of laboratory experiment time-series measurements of the cyanobacterium *Prochlorococcus*, Earth’s most abundant photosynthetic organisms, demonstrated the performance improvements of our approach over current models, and isolated previously ignored respiratory and exudative carbon losses as critical parameters for the modeling of microbial population dynamics.

## Introduction

Marine phytoplankton are photosynthetic microorganisms that account for up to half of global net primary production [1]. As such, the population dynamics of these organisms are crucial to understanding the global carbon cycle [2, 3]. One key aspect of phytoplankton populations is the growth rate, typically defined as the rate of increase in population biomass over time per unit of existing biomass. Direct *in-situ* measurement of this bulk quantity is obscured by heterotrophic biomass and detrital material, which constitute a variable fraction of the particulate organic carbon pool [4]. Several different methodologies have been employed to estimate *in-situ* phytoplankton growth rates; however, previous estimates relied on analytically challenging and low-throughput methods such as the radiometric turnover times of ^14^C labeled chlorophyll [5] and ^32^P labeled ATP [6], cell cycle analysis [7], and the dilution method [8]. While taxon-specific growth rates might be estimated with these methods, they often suffer from large uncertainties caused by coarse sample time resolution or experimental artifacts (collectively known as “bottle effects”; e.g., [9]). The emergence of continuous flow cytometry in ocean surveys [10–12] provides high resolution, taxon-specific measurements of the abundance and size of individual phytoplankton cells and offers a high-throughput *in-situ* alternative. In principle, measurements of cell abundance across different sizes over time provide a means to directly derive rates of carbon fixation and cell division [4], but the mechanistic modeling frameworks are currently underdeveloped and cannot accurately isolate these implicit rates from other cellular processes.

The class of mechanistic models we focus on consists of stage-structured matrix population models (MPMs), which estimate demographic rates from measurements of abundance across life-cycle stages [13], often defined by the age or size of individuals. For example, tree species produce seeds once they have reached a particular size [14] and fish species maximize reproduction at a critical age [15]. These models assume that individuals in a population can be classified into *m* discrete stages that define their response to the environment modeled as a discrete-time process. MPMs assume that the state of the population at time *t* + 1 can be written in terms of the state of the population at time *t* and a set of transition rates [16]:

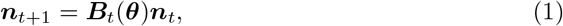

where ***B***_*t*_(***θ***) is a *projection matrix* that defines the possibly time-dependent population dynamics, ***θ*** is a parameter vector, and ***n***_*t*_ is a vector representing the number of individuals in each stage at time *t*, which defines the state of the population. The vector ***θ*** includes both biological and mechanistic parameters to model population dynamics and is the target of parameter estimation [17].

In recent years, size-structured MPMs have gained popularity for estimating rate parameters of phytoplankton populations by mechanistically describing changes in microbial cell size distributions over the day-night cycle [18–24]. For instance, MPMs have been employed to estimate daily division rates of the picocyanobacterium *Synechococcus* and picoeukaryotic phytoplankton based on a 13-year hourly time series from a coastal location in the Atlantic Ocean using a submersible flow cytometer [19, 23, 24]. In the North Pacific Subtropical Gyre, similar MPMs were used to estimate daily and hourly division rates of another picocyanobacterium, *Prochlorococcus*, based on continuous flow cytometry measurements taken over two research cruises [21].

In these studies, cell size measurements provided by high-frequency flow cytometry were used to define the life-cycle stages of the population. These models assumed that changes in the cell size distribution over the day-night cycle are driven only by two biological processes: 1) carbon fixation via photosynthesis and 2) cell division; other processes such as respiration and exudation, which lead to cell shrinkage, are omitted. In previous investigations, model performance was judged on the goodness of fit to the size distribution data rather than the plausibility of model parameters, in part due to the difficulty of directly assessing biological feasibility of demographic rates of microbial populations. Uncertainty quantification for model parameters typically involved refitting methods or was ignored entirely, omitting critical context from the inference procedure. As a consequence, these MPMs [18, 19, 21, 24] contain loosely constrained model parameters that can lead to transition matrices with biologically implausible estimates.

Here, we extend existing size-structured MPMs to test a set of underlying assumptions describing population dynamics over the day-night cycle and to improve parameter interpretability and model performance. Model estimates are computed using the Bayesian implementation in the probabilistic programming language Stan [25], through which we provide statistically rigorous parameter uncertainty intervals while constraining parameter values by incorporating prior scientific knowledge. This approach enabled an evaluation of the sensitivity of posterior distributions to sampling size, sampling frequency, and initial conditions. In the following, we test nine MPMs that differ in their parameterizations of three transition terms: cell division, carbon fixation, and carbon loss (Fig 1) which describe the dynamics of the picocyanobacterium *Prochlorococcus*, Earth’s most abundant phytoplankton [26].

**Fig 1.**
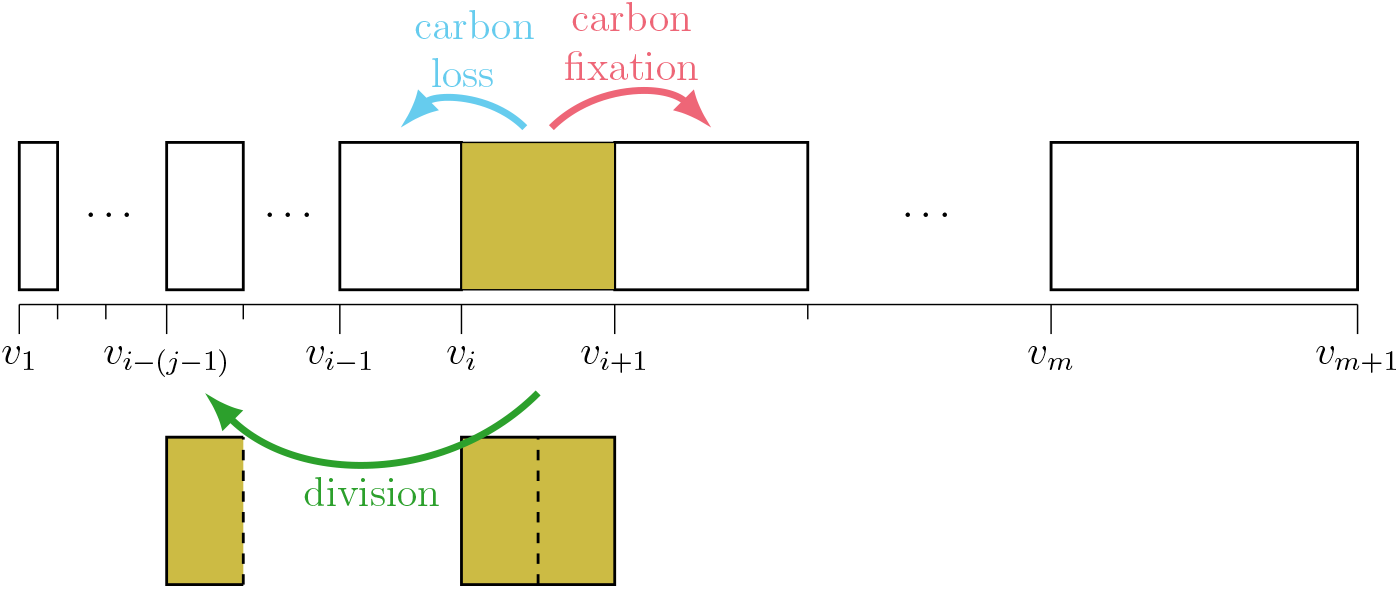
MPM size classes and transitions. Schematic of the MPM’s cell size classes and its three class transitions: carbon fixation, division, and carbon loss. The boundaries of the *m* cell size classes (*v_i_* for *i* = 1, 2, … , *m* + 1) are logarithmically spaced, so that cells can transition to a size class that is exactly half their original size when they divide. For this purpose, the integer *j* is selected so that 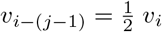 for *i* > *j*; cells in the first *j* size classes cannot divide.

We evaluated the performance of our models on a publicly available dataset of laboratory experiment time-series measurements of a high-light adapted strain of *Prochlorococcus* [27] collected during the exponential phase of batch growth over two simulated day-night cycles (Fig 2). This dataset contains cell size distributions derived from flow cytometry (Fig 2 A, B), cell abundance and light measurements (Fig 2 C), and measurements of carbon fixation, carbon loss, and division (Fig 2 D) at two-hour intervals. Division rates are derived from changes in cell abundances while carbon loss is estimated from other measurements (see Experimental data below). We fit our models to the size distribution data (Fig 2 A, B) and then evaluated how well each model was able to reproduce the observed parameters at daily and hourly time scales. All models used a logarithmically-spaced discrete cell size distribution, permitting cells to divide into two daughter cells that are half their size (Fig 1). While our simplest model has no size-dependence for carbon fixation and lacks a carbon loss term, the more complex models include size-dependence for all three transitions, explained below. Finally, we converted model parameters to estimates of biological rates such as carbon fixation and carbon loss, connecting microbial growth processes to the marine carbon cycle.

**Fig 2.**
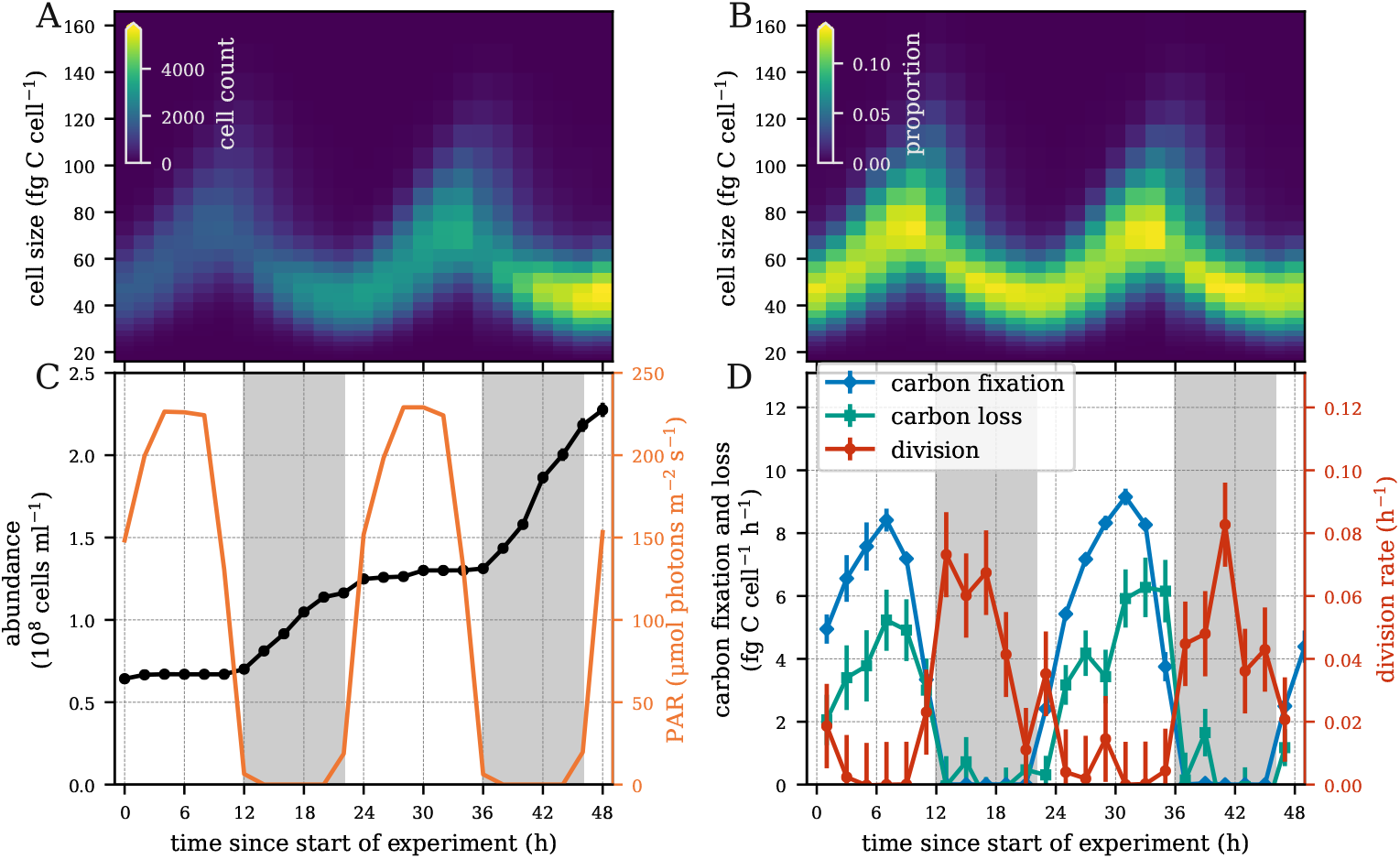
Laboratory *Prochlorococcus* time series measurements. (A) Heatmap of the number of cells and (B) relative cell abundances in each size class measured every two hours over a 48-hour period. (C) Cell abundance and photosynthetically active radiation (PAR). (D) Hourly carbon fixation, carbon loss, and division rates. Error bars indicate one standard deviation based on two technical and two biological replicates.

## Results

### Models

Past work has assumed that changes in cell size result from two processes: carbon fixation and cell division [18–24]. We built upon these studies by evaluating the relevance of a range of assumptions and testing models that include an additional process: cell shrinkage through exudation and respiration. Another assumption of past models is that division is a monotonically increasing function of size, i.e. larger cells are more likely to divide than smaller cells. This implies that the highest rate of cell division should occur when cells reach their largest size. However, the peak of average cell size in *Synechococcus* and *Prochlorococcus* occurs during daylight while the peak of division usually occurs at night [28]. In the *Prochlorococcus* culture dataset used in our work, hourly cell division lagged 4-8 hours behind the peak of cell size (Fig 3 A). In fact, hourly division rates showed little correlation with mean cell size (Fig 3 B). When comparing the size distribution at 13 hours (peak in cell division) and at 35 hours (almost no division) after the start of the experiment, we see that the size distributions are fairly similar despite the large difference in division rates (Fig 3 C). However, we observed a strong correlation (r=0.84) between hourly division rate and mean cell size with a 6-hour lag (Fig 3 D), suggesting that cell division is dependent on cell size as well as additional processes. For instance, cell division in photosynthetic organisms is tightly regulated by light, although the onset of the cell cycle in *Prochlorococcus* does not seem to be strictly light dependent [30]. We therefore tested two different parameterizations for estimating cell division. In the first, cell division is constrained to be a monotone function of cell size, but constant over time, as in previous studies. In the second, cell division still increases monotonically with cell size but is allowed to vary over time. We also considered size dependence in carbon fixation through power-law relationships supported by experimental evidence [29]. Finally, we implemented a “free” parameterization in which carbon fixation and carbon loss rates are estimated separately for each size class, in order to provide enough flexibility for the model to capture biological processes that are not explicitly accounted for in our models.

**Fig 3.**
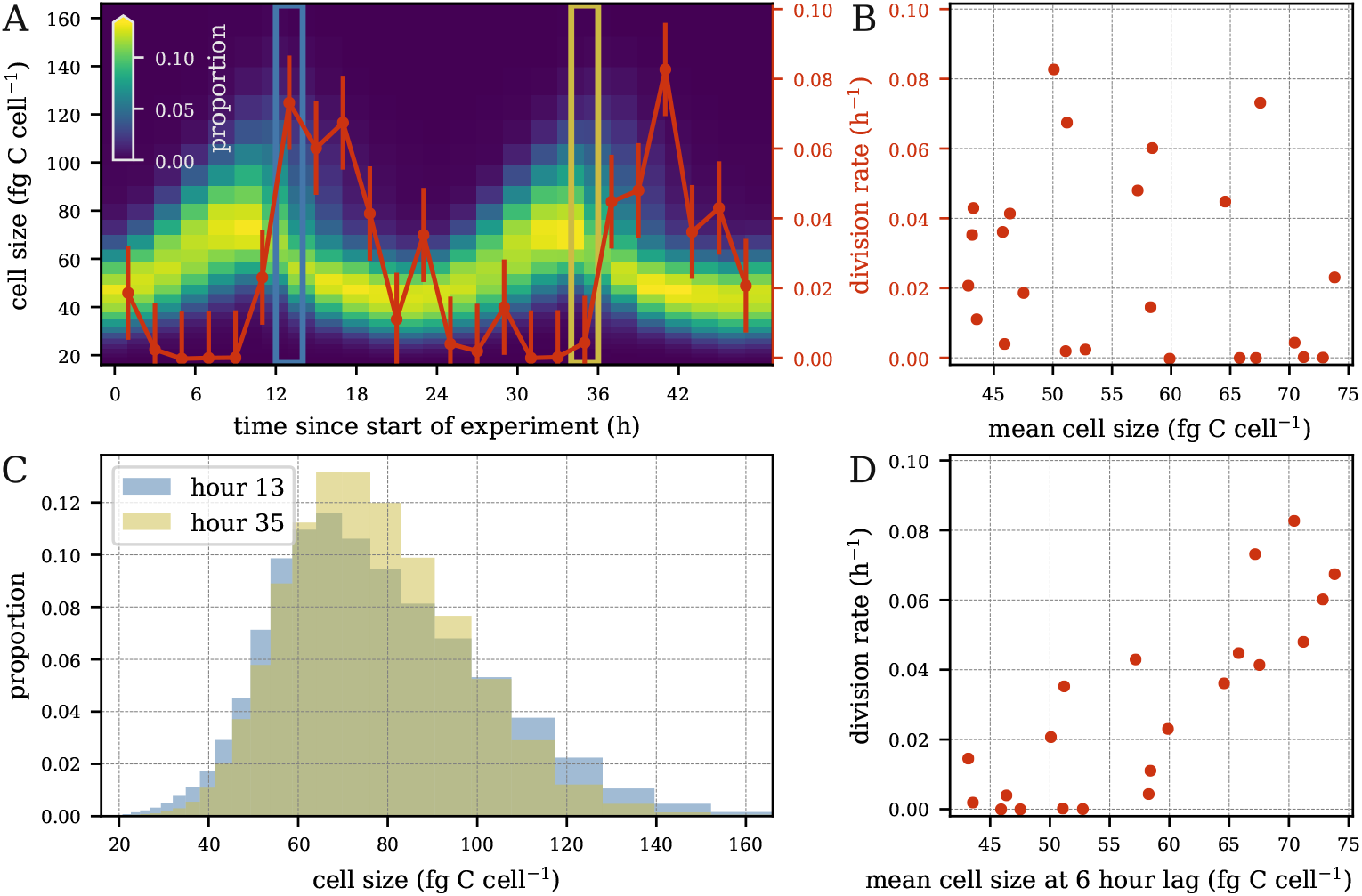
Hourly division rates vs. average cell size. (A) Phytoplankton size distribution overlaid with hourly division rates (red curve; error bars indicate one standard deviation based on two technical and two biological replicates). Division rate and size distribution at *t* = 13 (blue box) and *t* = 35 (gold box). (B) Hourly division rates vs. mean cell size. (C) Cell size distribution at time *t* = 13 (blue) and *t* = 35 (gold). (D) hourly division rate at time *t* vs. mean cell size at time *t* − 6.

We distilled our assumptions into a set of 9 models of differing parameterizations (Table 1). Each model is identified by a subscript consisting of three letters corresponding to the parameterizations of carbon fixation, division, and carbon loss, respectively. The first letter in each model name corresponds to the carbon fixation parameterization. The letter **b** in carbon fixation indicates a basic parameterization in which carbon fixation is assumed to be constant as a function of size. The letter **p** indicates a power-law relationship with respect to size and **f** represents a free parameterization where each size class may have its own rate of carbon fixation. With respect to division, represented by the second letter of the model name, the letter **m** indicates a monotone division rate as a function of size with no time-dependence, while **t** indicates a parameterization that also includes time-dependence in division. The third letter, indicating the carbon loss parameterization, can be **b** (basic) or **f** (free parameterization) as in carbon fixation, or **x** for a model with no carbon loss. As an example, we refer to our simplest model as *m*_bmx_, denoting that it has basic carbon fixation without size-dependence, division rates that monotonically increase with cell size, and no carbon loss term.

**Table 1.** Key models.

| Model* | Growth | Division | Loss |
| --- | --- | --- | --- |
| $m_{\text{bm}x}$ | basic | monotonic | x (no loss) |
| $m_{\text{bm}b}$ | basic | monotonic | basic |
| $m_{\text{pmb}}$ | power-law size-dependence | monotonic | basic |
| $m_{\text{fmb}}$ | free size-dependence | monotonic | basic |
| $m_{\text{fmf}}$ | free size-dependence | monotonic | free size-dependence |
| $m_{\text{btb}}$ | basic | time-dependent | basic |
| $m_{\text{ptb}}$ | power-law size-dependence | time-dependent | basic |
| $m_{\text{ftb}}$ | free size-dependence | time-dependent | basic |
| $m_{\text{ftf}}$ | free size-dependence | time-dependent | free size-dependence |

\*The letters in the subscript of the model name denote the growth, division, and loss parameterizations used in the model, respectively.

The two division parameterizations split our models into two groups. Within each group, models contain more parameters down the rows of Table 1. Between the two groups, models with time-dependent division contain more parameters than their time-independent versions. Thus, model *m*_bmx_ was the simplest model and most closely represented previous MPMs applied to microbial communities, while model *m*_ftf_ is the most complex with respect to the number of parameters.

We fit these 9 models to a dataset gathered in a laboratory experiment. Rates of division, carbon fixation, and carbon loss were estimated on both daily and hourly timescales. In the following section, we examine daily rate estimates, which have been the primary target of inference in past work. Then, we further assess the model rate estimates at an hourly timescale to inspect the behavior of our models within diel cycles. Furthermore, we explore the relationship between cell size and division, carbon fixation, and carbon loss. Finally, we examine the relationships between the estimated parameter values and perform observation sensitivity experiments.

### Estimation of daily rates

We first assessed our models’ ability to recreate the observed *Prochlorococcus* cell size distribution. Then, we examined whether an improved fit to the size distribution data resulted in improved model performance by comparing model estimates of daily average carbon fixation, carbon loss, and division rates to independent measurements from laboratory data. Finally, we investigated model estimated photosynthetic parameters.

As expected, the MSE of the predicted cell size distribution decreased as the number of model parameters increased (Fig 4A). Critically, however, this improved fit did not correlate with better daily rate estimates. One of the most important parameters estimated by the models is the daily rate of cell division, see Eq (4). The observed daily division rate in the population was 0.63 ± 0.01 d^−1^. However, the simplest model *m*_bmx_ overestimated this rate by nearly a factor of two (Fig 4 B; 1.06 ± 0.05 d^−1^). This may stem from the fact that this model did not include carbon loss; thus, it attributed any reduction in cell size to cell division. Model *m*_bmb_, which adds respiratory/exudative carbon loss, was able to accurately estimate the daily division rate (0.63 ± 0.02 d^−1^), while all other models produced less accurate estimates, despite lower MSE of the predicted cell size distribution.

**Fig 4.**
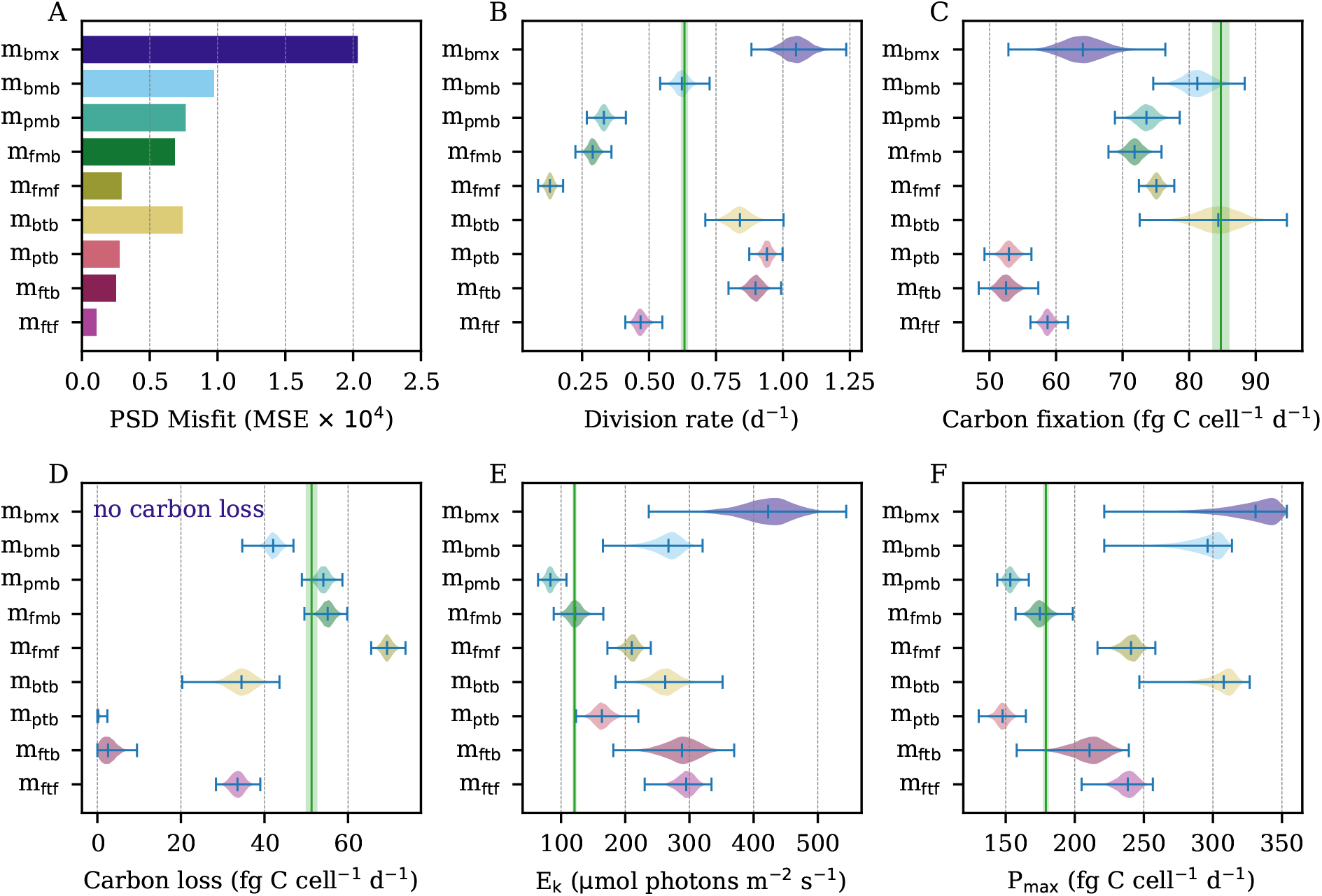
Model predicted daily rate parameters. (A) Mean squared error (MSE) of predicted proportions to the observed particle size distribution (PSD). (B) Predicted daily division rates. (C) Predicted daily carbon fixation. (D) Predicted daily carbon loss. (E) Predicted photosynthetic saturation parameter. (F) Predicted maximum photosynthetic rate. (B-F) Green vertical lines indicate ground truth calculated from data. Green shaded areas indicate uncertainty surrounding ground truth measurements. Model estimates shown as posterior distributions.

Model *m*_bmb_ also performed well in estimating daily rates of carbon fixation and loss (Fig 4 C,D). Again, the models with the best fit to the size distribution (*m*_fmf_, *m*_ptb_, *m*_ftb_, *m*_ftf_) exhibited lower accuracy in their estimates of these rates. Interestingly, the addition of size-dependent carbon fixation (*m*_pmb_, *m*_fmb_) resulted in underestimation of daily carbon fixation (75.57 ± 1.00 fg C cell^−1^ d^−1^ and 73.77 ± 1.00 fg C cell^−1^ d^−1^ for *m*_pmb_, *m*_fmb_, respectively) and cell division (0.33 ± 0.02 d^−1^ and 0.29 ± 0.02 d^−1^, respectively) but improved estimates of daily carbon loss. The further addition of size-dependence in carbon loss (*m*_fmf_) led to overestimates of daily carbon loss and even lower division rate estimates, indicating that this model attributes too much of the observed decreases in cell size to carbon loss rather than cell division. Other than *m*_btb_, which exhibits more instability than other models and whose results may therefore not be reliable (see Observation sensitivity experiments section), models that added time-dependent division (*m*_ptb_, *m*_ftb_, *m*_ftf_) greatly underestimated both carbon fixation and carbon loss rates. Models without size-dependent carbon loss (*m*_ptb_, *m*_ftb_) estimated essentially no carbon loss, leading to inflated division rates as nearly all cell size decreases were attributed to cell division. This effect was counteracted to some extent by the inclusion of size-dependent carbon loss (*m*_ftf_), although both the daily division rate and carbon fixation were underestimated.

Finally, we examined the photosynthetic saturation parameter *E_k_* and the maximum light-saturated photosynthetic rate P_max_, two components of the mechanics of carbon fixation (see Carbon fixation section). Model *m*_bmx_ shows the worst performance for these parameters, but *m*_bmb_ also greatly overestimates both quantities despite accurate estimation of daily carbon fixation, highlighting potential identifiability issues - i.e. similar daily carbon fixation rates can be obtained by different means, as carbon fixation decreases with higher values of *E_k_* but increases with higher values of P_max_. Interestingly, *m*_pmb_ and *m*_fmb_ had much more accurate estimates of the photosynthetic parameters, despite lower accuracy in overall daily carbon fixation. Size-dependent carbon loss (*m*_fmf_) and time-dependent division (*m*_btb_, *m*_ptb_, *m*_ftb_, *m*_ftf_) resulted in poorer estimates of the photosynthetic parameters relative to *m*_fmb_.

Overall, the simplest model *m*_bmx_ showed the poorest performance in estimation for nearly every category, highlighting the importance of accounting for carbon loss in our models. There is no model that performed best with respect to all of the daily rate estimates we included in our tests; *m*_bmb_ created the best division and carbon fixation estimates, *m*_fmb_ provided the best performance for *E_k_* and P_max_, and *m*_pmb_ most accurately predicted daily carbon loss.

### Estimation of hourly rates

In addition to the analysis of daily rate parameters, we examined the models’ abilities to recreate population dynamics at hourly resolution (Fig 5) to determine whether discrepancies between model predictions and observations occur at a particular time of the diel cycle and to help us identify the relevant biological processes at play. For clarity, we show here the results of the five most distinct models (*m*_bmx_, *m*_bmb_, *m*_fmf_, *m*_ptb_, and *m*_ftf_); results for all nine models can be found in the SI (Fig S8). While some of our models were able to estimate the daily rates of cell division, carbon fixation, and carbon loss accurately, the hourly patterns were more difficult to replicate (Fig 5A-C). As expected by the relationship between cell size and hourly division rates (Fig 3), models that assume that cell division is only size-dependent (*m*_bmx_, *m*_bmb_, *m*_fmf_) predicted the timing of cell division to be 4 to 8 hours too early (Fig 5A). On the other hand, models with both time-dependent division and size-dependent carbon fixation (*m*_ptb_, *m*_ftf_) were able to more accurately predict the timing of cell division. However, these models either overestimated division during the morning (*m*_ptb_) or underestimated division at dusk (*m*_ftf_), thus leading to the inaccurate daily rates as discussed above. All models were able to capture the timing of carbon fixation, which is tied to the amount of incident light (Fig 5 B). However, most models tended to underestimate the amount of fixed carbon, with *m*_bmb_ coming closest to capturing the dynamics observed in the data. Surprisingly, the timing of carbon loss computed from the data (Fig 5 C) closely matched that of carbon fixation. Our models tended to underestimate carbon loss during daytime peaks and overestimate it at night.

**Fig 5.**
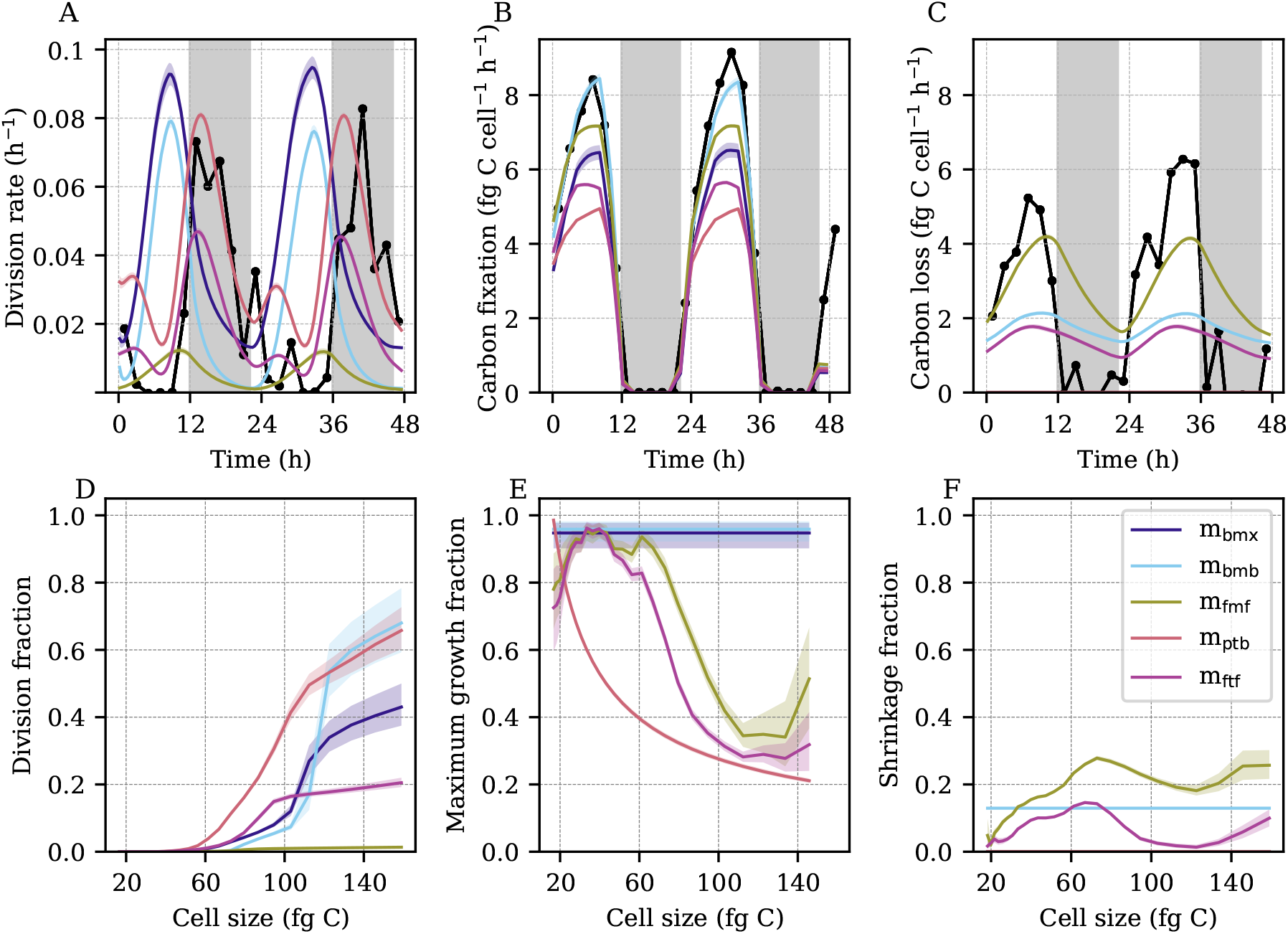
Model predicted hourly rate parameters. (A) Observed (black) and predicted (colored bands) hourly division rates. (B) Observed (black) and predicted (colored bands) hourly carbon fixation. (C) Observed (black) and predicted (colored bands) hourly carbon loss. (A-C) Black points indicate ground truth calculated from data. (D) Predicted cell division fraction as a function of cell size. (E) Predicted light-saturated cell growth (carbon fixation) fraction as a function of cell size. (F) Predicted cell shrinkage (carbon loss) fraction as a function of cell size. (A-F) Colored bands indicate model estimates. Shading indicates the first to third quartiles of the posterior distributions. (D-F) Fractions correspond to MPM transitions over a 20-minute time period.

To further explore the predicted dynamics of division, carbon fixation, and carbon loss, we investigated the predicted proportions of cells undergoing each of these transitions as a function of cell size (Fig 5 D-F). The estimated shape of the size-division relationship tended to follow a sigmoidal pattern for all models: the fraction of dividing cells increases sharply above a critical size, which varied from 60 to 110 fg C depending on the model (Fig 5D). We note that the model that best estimated the daily division rate (*m*_bmb_) predicted cell division to occur mostly in the largest size classes (> 110 fg C), which resulted in accurate amplitudes of hourly cell division rates, albeit at a 6-hour phase shift. In general, models that overestimated cell division rates (*m*_ptb_) predicted higher proportions of dividing cells for smaller sizes, while models that underestimated division (*m*_fmf_, *m*_ftf_) estimated smaller proportions of dividing cells within the larger size classes. The exception to this trend is *m*_bmx_, which generally estimates a comparable or lower division fraction than *m*_bmb_ at a given size yet overestimates cell division. Because *m*_bmx_ contains no carbon loss, it predicts more large cells to be present in the distribution, hence increasing the predicted division rate relative to *m*_bmb_ even if the division fraction is lower.

Meanwhile, model estimates of the size-dependence of carbon fixation generally estimated high values for the peak maximum growth fraction (Fig 5 E). Models that assumed constant maximum growth (*m*_bmx_, *m*_bmb_) estimated this fraction to be near one. Interestingly, models with a free parameterization of size-dependent carbon fixation (*m*_fmf_, *m*_ftf_) generally predicted larger cells to have a lower maximum growth fraction, as in the power-law formulation (*m*_ptb_). The predicted fractions of cell shrinkage tended to be significantly lower than the fractions of maximum growth, ranging from negligible to about one-fifth of the peak maximum growth fraction (Fig 5E, F). In the two models with size-dependent carbon loss rates (*m*_fmf_, *m*_ftf_), the predicted fraction of cell shrinkage generally increased with cell size. However, both models estimated a sharp drop near the same critical sizes at which the division fraction sharply rose, suggesting that the models assign the decreases in cell size to cell division rather than carbon loss for larger but not smaller cells. These results suggest a trade-off of daily and hourly rate estimates between our models: models that produced some of the most accurate daily estimates of cell division, carbon fixation, and carbon loss showed a systematic offset in timing of cell division, while the models which accurately captured the timing often performed less well in estimating the daily average rate.

### Posterior parameter distributions

As the cell size distribution is used for model fitting, a model may be able to accurately capture the net effect of the parameters despite failing to accurately capture the value of each parameter individually, highlighting potential identifiability issues. We therefore examined the bivariate joint posterior distributions of estimated rates of daily cell division, carbon fixation, and carbon loss as well as photosynthetic parameters to better understand the mechanics of the MPMs and the interdependencies of their parameters. We focused on two models: *m*_bmb_, which had the best overall performance on daily rates of cell division, carbon fixation, and carbon loss but failed to predict the timing of cell division, and *m*_ftf_, which was best able to predict the timing of cell division but failed to provide accurate daily rates (Fig 6). A strong correlation between daily carbon fixation and carbon loss was observed in the posterior distributions of both models (r = 0.61 and 0.81 for *m*_bmb_ and *m*_ftf_, respectively; Fig 6 J,K), which was expected since the carbon fixed by photosynthesis fuels respiration and exudation. However, the relationship between carbon fixation and cell division differed between the two models (Fig 6 F,O). Carbon fixation and cell division were positively correlated (r = 0.64) in *m*_bmb_, which makes intuitive sense since the faster the cells grow, the faster they divide (Fig 6 F), while a negative correlation (r = −0.46) was observed in *m*_ftf_ (Fig 6 O). This negative relationship likely stems from the fact that daily division rate and carbon loss in *m*_ftf_ were strongly negatively correlated (r = −0.88, Fig 6 L), while this relationship was much weaker in *m*_bmb_ (r = −0.19, Fig 6I). As carbon fixation and carbon loss are tightly correlated, carbon loss may mediate the observed negative relationship between carbon fixation and daily division in *m*_ftf_, making it more difficult for this model to disentangle these two processes than in *m*_bmb_.

**Fig 6.**
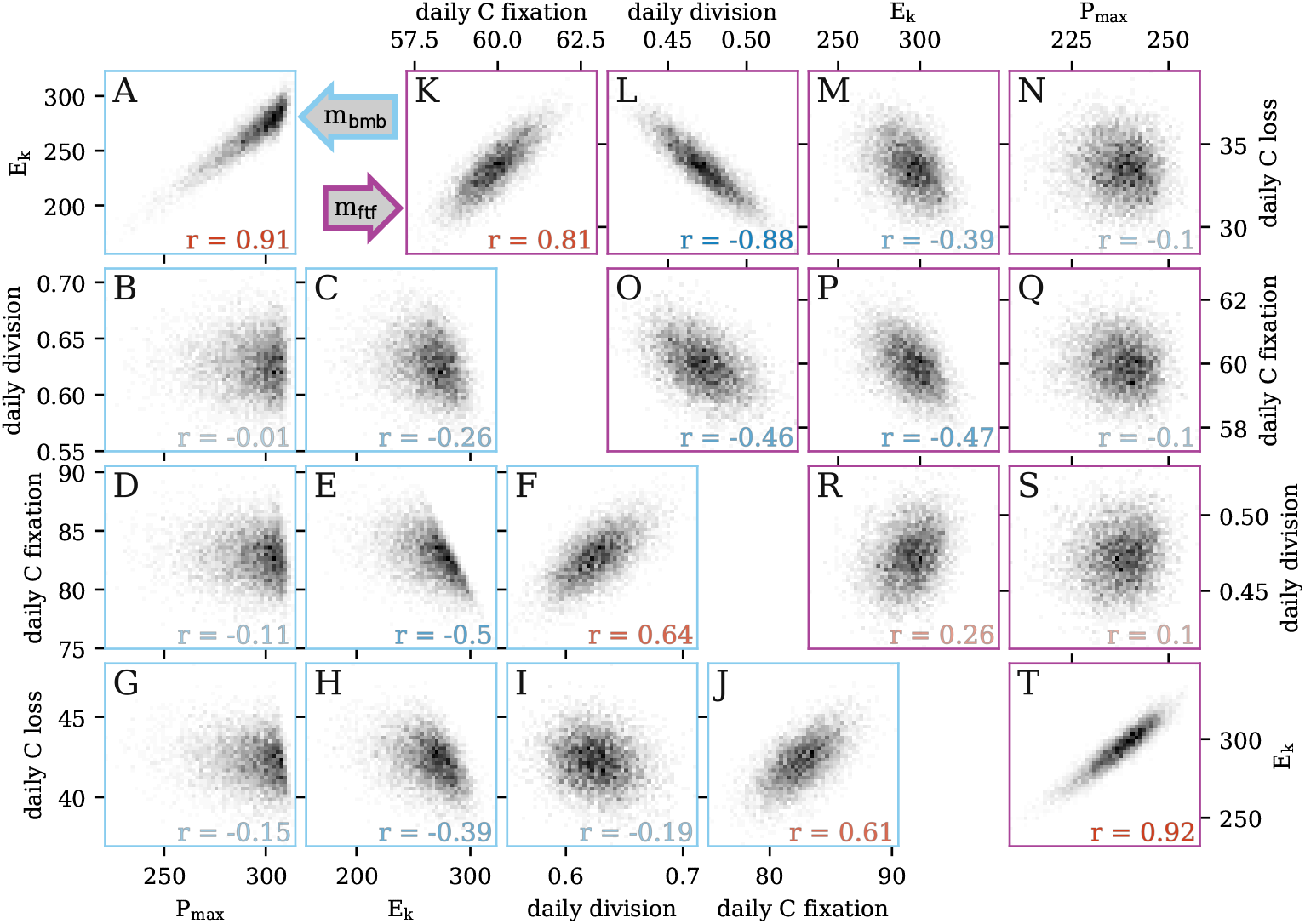
Bivariate posterior distributions. Scatter plots of the bivariate posterior distributions of select parameters for the models (A-J) *m*_bmb_ and (K-T) *m*_ftf_.

The shape of the posterior distribution highlights the strong relationship between P_max_ and *E_k_* (Fig 6 A,T); increases in P_max_ and reduction of *E_k_* both increase carbon fixation in different ways (see Eq (12)), which would explain why *m*_bmb_ could accurately estimate daily carbon fixation albeit with inaccurate estimates of photosynthetic parameters. The strong dependence structure between parameters shows that it is important to consider the joint distributions of the parameters and not solely focus on the marginal posterior distribution for each parameter. It also demonstrates that the size-distribution data itself cannot constrain all parameters, emphasizing the importance of prior knowledge and the prior distribution for limiting the parameter distributions.

### Observation sensitivity experiments

In order to quantify the impact of changes in the size distribution data on model parameter estimates, we performed two sets of experiments. In the first, we used a sliding window approach to assess the effect of changing the start time of the 48-hour time series on model output. In the second, we studied the robustness of the models to changes in the sampling resolution of observations.

In the sliding window experiment, we extended the normalized size distribution time series by appending the data to itself, thereby creating a four-day dataset. Then, we estimated parameters and initial conditions within a 48-hour window that was moved forward in time in four-hour increments. Details about the setup of the sliding window experiments and their results can be found in the SI (Section S1). With the exception of *m*_bmx_ and *m*_btb_, all models exhibited a high degree of stability in their estimates for each window, indicating that the starting time of the model fitting procedure had a very limited effect on the models’ parameter estimates. Some deviations were however noticeable when the window start time was near the peak of the cell size distribution, at which the difference between observations and model predictions is most pronounced. For *m*_bmx_ and *m*_btb_, estimates showed a high degree of variability among windows, suggesting that the results of these models may not be as stable or reliable as the others.

In the second set of experiments, we performed holdout validation experiments in which time points of the size distribution data were withheld from the training data used for model fitting. This holdout data was then used as a testing set and we computed the error for both datasets in order to examine our models’ out-of-sample performance and the stability of the parameter estimation relative to the full dataset. We conducted three experiments, sequentially removing an increasing amount of equally spaced data, roughly mimicking lab experiments in which measurements were collected at lower resolution. This procedure ensured that the daily cycle was sampled well and both days are represented equally in the training data. More details of this analysis can be found in the SI (Section S2). We found that parameter estimates and the observed cell size distribution remained stable when up to half of the data was removed from training, but out-of-sample performance deteriorated and parameter estimates differed significantly from those computed from the full data when two-thirds of the data was removed. This result suggests that our model could be applied to time series data collected at 4 hour interval and still provide accurate estimated daily rates of cell division, carbon fixation, and carbon loss.

## Discussion

In this work, we developed a flexible framework that allowed us to test multiple hypotheses regarding key biological processes that dictate phytoplankton cell growth, shrinkage, and division. Our investigation focused on a laboratory culture of the picocyanobacterium *Prochlorococcus*, whose dynamics over the diel cycle have been extensively studied [27]. We developed nine models that differed in their parameterizations of changes in cell size. In addition to a size-dependent relationship for cell growth and time-dependence in cell division, we considered respiratory and exudative carbon loss in our models, which had previously been neglected in similar studies [18–24]. To this end, we implemented our models within a Bayesian framework, which permitted us to incorporate prior information into the analysis to regularize parameter inference and avoid biologically implausible parameter values.

Herein, we showed that size-structured MPMs can be used to estimate not only rates of cell division but also carbon fluxes, thereby connecting microbial growth processes to the carbon cycle. The addition of carbon loss, which allows cells to shrink in size through a process other than cell division, improved the accuracy of model estimates and the fit to the size distribution data, with *m*_bmb_ successfully recovering the measured daily rates of cell division, carbon fixation, and carbon loss (Fig 4 B-D). More complex models, such as those with size-dependent carbon fixation and time-dependent cell division, provided better fits to the cell size distribution and photosynthetic parameter estimates but showed worse model performance in recovering the observed daily rate parameter values. This result indicates that model fit to the observed cell size distribution cannot be used as a proxy for overall model performance, as done in previous studies [18–24].

As expected from the lack of correlation between mean cell size and hourly division rate in the laboratory data (Fig 3), most of our models consistently predicted the peak of cell division about 4-8 hours earlier than observed in the data (Fig 5; Fig S8). This offset stemmed from the assumption that cell division (i.e. the separation of a single cell into two daughter cells) occurs instantaneously once the cells reach a certain size. While this assumption may be reasonable on daily time scales, it becomes problematic at hourly resolution; cell division is a complex process involving many components, each highly regulated to ensure that the separation of the cell into two daughter cells occurs only once DNA synthesis is completed, which takes between 4 and 6 hours depending on the strain and culture conditions [27, 30]. Here, the peak of DNA synthesis coinciding with the peak of cell size [27] suggests that cell size dictates the onset of DNA replication rather than the final separation of the cell into two daughter cells. Due to their greater flexibility, models with time-dependent division and size-dependent carbon fixation successfully captured the timing of cell division but failed to predict accurate rate estimates. Interestingly, models with a free parameterization of the size-dependent carbon fixation (*m*_fmb_, *m*_fmf_, *m*_ftb_, and *m*_ftf_) estimated less carbon fixation and more carbon loss in the large size classes which contains a large fraction of dividing cells (Fig 5 E,F; Fig S8 E, F). This result suggests that dividing and non-dividing *Prochlorococcus* cells may have a different carbon metabolism, as observed in other organisms [31].

Finally, we consider potential future directions for this work. One of the most interesting results in this study is the offset in the predicted and observed timing of division for the models with the most accurate daily division rate estimates. While the addition of time- and size-dependencies for cell division, carbon fixation and loss allowed our more complex models to capture the timing of cell division, their estimates of the magnitude of division and other rate parameters suffered. As stated above, we hypothesize that carbon metabolism differs between dividing and non-dividing cells, yet our current modeling framework requires extension of the stage structure to encapsulate this information in order to test such a hypothesis. A hybrid age- and size-structured MPM may therefore be better suited to assess the importance of including cell division duration to more accurately simulate the timing of *Prochlorococcus* division.

An exciting future extension of this work is application to an *in-situ Prochlorococcus* and *Synechococcus* dataset obtained from shipboard flow cytometers [32]. Additional processes not accounted for in this study, such as grazing and viral lysis, which could potentially affect phytoplankton size distributions, will need to be tested. The flexibility of our modeling framework provides a valuable basis for examining and evaluating MPM results in the face of more complex datasets, which could further improve our understanding of the dynamics of marine microorganisms and their contributions to the carbon cycle.

## Materials and methods

### Microbial MPM

The aim of the MPM applied to microbial populations is to mechanistically describe the evolution of the population over a day/night cycle. Typically, the target of inference is the daily division rate, which cannot be measured directly from changes in cell abundance measured in the field due to cell mortality caused by grazing and viral lysis as well as physical processes that can add or remove cells from the sampled population. Thus, in order to estimate this quantity, we infer it via observed changes in the relative abundance distribution over time. Past work has accomplished this by focusing on modeling two cellular processes: cell division and carbon fixation; in this work, we additionally consider carbon loss. We tested nine MPMs involving these processes that varied in their complexity. All inference was carried out using the Bayesian modeling software Stan, see Implementation section below.

### Preliminaries

The MPM operates on discrete scales in both cell size and time. Therefore, there are two user-defined discretization parameters: Δ*v* ∈ ℝ^+^ is the size discretization parameter and *dt* ∈ ℝ^+^ is the time discretization parameter in hours. We choose the former such that Δ*v*^−1^ ∈ ℕ so that division corresponds to shifting 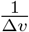 size classes, see (2). We choose the latter to match our observation resolution; as the dataset has observations every 2 hours, we enforce *dt*^−1^ ∈ ℕ. In addition, we define *m* ∈ ℕ the total number of discrete size classes and *v*_1_ the minimum possible cell size, to define *m* + 1 size class boundaries:

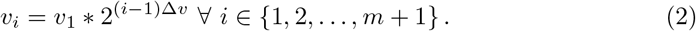

If a cell is of size *x* where *v_i_* ≤ *x* < *v*_*i*+1_, then the cell belongs to size class *i*.

Furthermore, we denote 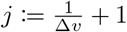 so that *v_j_* = 2*v*_1_, i.e. only cells of size class *j* or greater can undergo cell division, see (7). For size-dependent parameterizations (see (12)), we treat cells in size class *i* as having size

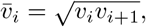

that is, they are assigned the geometric mean of the size class boundaries. In this work, we set *m* = 27, 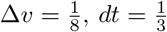 hour, and *v*_1_ = 16 fg C.

### Model inputs

The observations 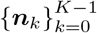 consist of cell counts across the *m* discrete size classes at *K* ∈ ℕ time points; that is, ***n***_*k*_ ∈ ℕ^*m*^ ∀ *k* ∈ {0, 1, 2, … , *K* − 1}. We denote the set of observation times as 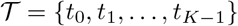, where *t_k_* ∈ ℕ refers to the time in hours of the *k*^th^ observation. For each *k*, we also define the simplex 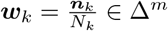, where 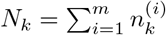 is the total number of cells observed at time *t_k_*. Observations also include measurements of photosynthetically active radiation (PAR), interpolated at the times 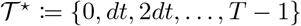, where the times are in hours; this information is used to estimate carbon fixation. We denote these values as 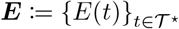. In our case, we have *T* = 47, *K* = 24, and 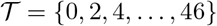.

### Model output

Microbial MPMs make projections operate differently from the formulation in (1). The predicted counts are normalized at each time step so that model projections estimate the relative abundance:

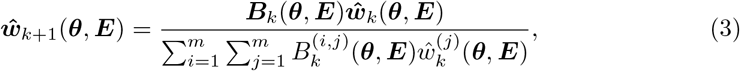

where ***θ*** is a parameter vector and ***B***_K_ (***θ, E***) ∈ ℝ^*m*×*m*^ is a projection matrix depending on model parameters, time, and incident light, see the Projection matrix section below. This formulation does not use the counts to estimate division rate directly, allowing for valid estimates even when mortality and physical movement of cells occur, so long as these processes do not affect the relative size distribution. We estimate the posterior distributions of the model parameters from their prior distributions and the likelihood of the data 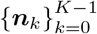 given the parameters (see Model likelihood section). The primary goal of inference is the daily division rate *μ*, defined as the exponential growth constant:

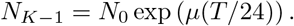

Recall that *T* = *t*_*K*−1_ is the time of the last observation in hours; thus, *T/*24 is the length of the time series in days. Rearranging the above equation, we obtain

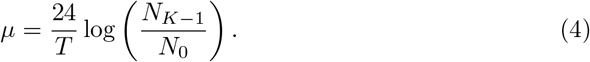

As populations in their natural environments undergo cell loss due to cell mortality (due to grazing and viral lysis) and physical processes that can add or remove cells, a normalization step (3) was applied to estimate division rate based on relative cell abundance, as in past applications [18, 19, 21]. By removing the normalization step, we estimate the relative increase in cell number caused by cell division. We therefore obtain the following estimator for the division rate:

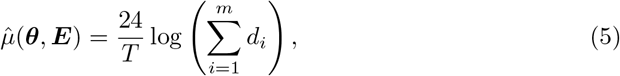

where ***d*** ∈ ℝ^*m*^ is defined as

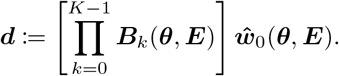

### Model likelihood

We use the following statistical model to assess the fit to the data:

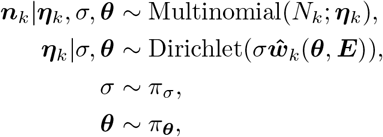

where *σ* is a real-valued concentration parameter, ***θ*** is a parameter vector, and *π*_·_ denotes the corresponding prior distributions (see Table 2). Thus, similar to [19], the model likelihood can be written as

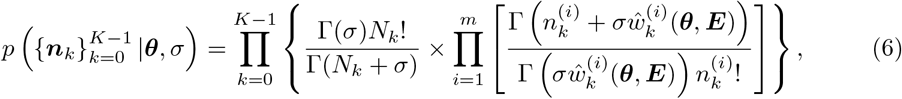

where 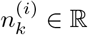 is the *i*^th^ entry of ***n***_*k*_ and 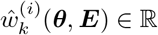 is the *i*^th^ entry of 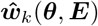. The posterior is proportional to the product of the likelihood and the prior distribution according to Bayes’ theorem; thus, we have

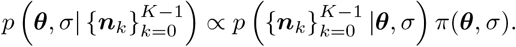

**Table 2.** List of model parameters.

| Name | Used in | Description | Units | Bounds | Prior |
| --- | --- | --- | --- | --- | --- |
| $w(0)$ | all models | initial conditions | – | simplex | Dirichlet($\mathbf{1}_m^T$) |
| $E_k$ | all models | light-dependent growth parameter | $\mu\text{mol photons m}^{-2} \text{ s}^{-1}$ | [0, 5000] | normal(1000, 1000) |
| $\delta_{\max}$ | all models | maximum division rate | $\text{d}^{-1}$ | $[0, \frac{1}{\Delta_t}]$ | uniform($0, \frac{1}{\Delta_t}$) |
| $\delta_{\text{incr}}^{(i)}$ | all models | increment in division rate, size class $i$ | – | [0, 1] | uniform(0, 1) |
| $\gamma_{\max}$ | all but $m_{f..}$ | maximum carbon fixation rate | $\text{d}^{-1}$ | $[0, \frac{1}{\Delta_t^*}]$ | normal(10.0, 10.0) |
| $\beta_\gamma$ | $m_{p..}$ | exponent in carbon fixation power law | – | [−10, 10] | normal(0, 0.1) |
| $\gamma_{\max}^{(i)}$ | $m_{f..}$ | maximum carbon fixation rate, size class $i$ | $\text{d}^{-1}$ | $[0, \frac{1}{\Delta_t^*}]$ | normal($\mu_\gamma, \sigma_\gamma$) |
| $\mu_\gamma$ | $m_{f..}$ | hierarchical prior for mean of $\gamma_{\max}^{(i)}$ | $\text{d}^{-1}$ | $[0, \frac{1}{\Delta_t^*}]$ | normal(10.0, 10.0) |
| $\sigma_\gamma$ | $m_{f..}$ | hierarchical prior for s.d. of $\gamma_{\max}^{(i)}$ | $\text{d}^{-1}$ | $[0, \infty[$ | exponential(0.1) |
| $\rho_{\max}$ | all but $m_{..f}$ | maximum carbon loss rate | $\text{d}^{-1}$ | $[0, \frac{1}{\Delta_t^*}]$ | normal(3.0, 10.0) |
| $\beta_\rho$ | $m_{..p}$ | exponent in carbon loss power law | – | [−10, 10] | normal(0, 0.1) |
| $\rho_{\max}^{(i)}$ | $m_{..f}$ | maximum carbon loss rate, size class $i$ | $\text{d}^{-1}$ | $[0, \frac{1}{\Delta_t^*}]$ | normal($\mu_\gamma, \sigma_\gamma$) |
| $\mu_\rho$ | $m_{..f}$ | hierarchical prior for mean of $\rho_{\max}^{(i)}$ | $\text{d}^{-1}$ | $[0, \frac{1}{\Delta_t^*}]$ | normal(10.0, 10.0) |
| $\sigma_\rho$ | $m_{..f}$ | hierarchical prior for s.d. of $\rho_{\max}^{(i)}$ | $\text{d}^{-1}$ | $[0, \infty[$ | exponential(0.1) |
| $\tau_{\text{control}}^{(i)}$ | $m_{.t.}$ | control point $i$ for time-dep. division spline | – | [0, 1] | beta(9, 1) |

Now, we characterize the parameter vector ***θ*** and the projection matrices ***B***_*k*_(***θ, E***), which generate model predictions.

### Projection matrix

The projection matrices 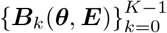 define the dynamics of the microbial population through three cellular processes: cell division, carbon fixation, and carbon loss. We assume that any individual cell can only undergo one of these three processes in each *dt* time step (it may also remain in the same size class). Thus, for each *k*, we first construct a set of matrices 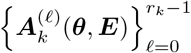, where *r_k_* ≔ (*t*_*k*+1_ − *t*_*k*_)*dt*^−1^ is the number of *dt* time steps between time *t_k_* and time *t*_*k*+1_. Once these matrices are defined, we have for each *k*:

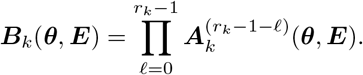

Each matrix 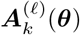 projects the process from time 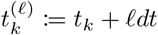 to time 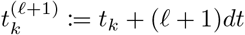.

Let *δ_i_*(*t*) ∈ [0, 1] denote the proportion of cells in size class *i* that divide in one *dt* time step at time *t*, *ρ_i_* ∈ [0, 1] the proportion of cells in size class *i* that shrink one size class in one *dt* time step given that they do not divide, and *γ_i_*(*t*) ∈ [0, 1] the proportion of cells in size class *i* that grow one size class in one *dt* time step at time *t* given that they neither divide nor shrink. Then, recalling that *j* denotes the index of the smallest size class which can undergo division, the entries of each matrix 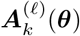 are defined as follows:

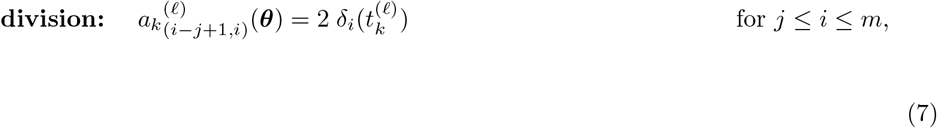

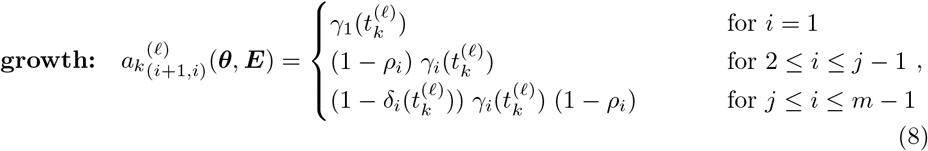

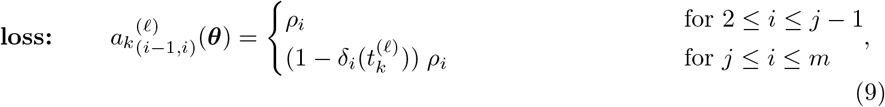

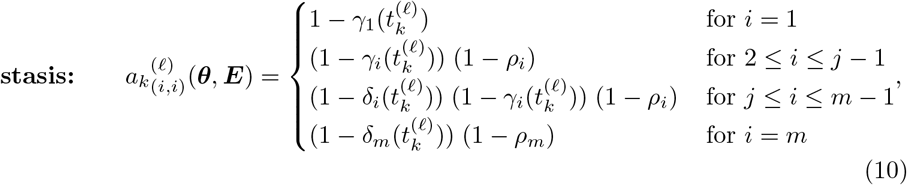

where again 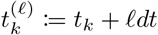. Here, only cell growth and stasis involve the PAR measurements ***E***. The coefficient 2 in equation (7) reflects the fact that when a cell divides, it creates two daughter cells. This is the reason the normalization step (3) is needed to maintain the sum-to-one constraint and also the reason (5), which omits the normalization, is able to estimate the rate of cell division.

### Parameterizations

In this work, we tested nine different microbial MPM’s. These models differed in their parameterizations of the three key processes we aim to quantify: cell division, carbon fixation, and carbon loss. Our most complicated models allow these processes to vary as functions of both time and cell size. The parameter vector ***θ*** controls the dynamics of these processes, while the concentration parameter *σ* allows for overdispersion in the data. We can divide the parameter vector ***θ*** into four components ***θ*** = (***θ***_*δ*_, ***θ***_*γ*_, ***θ***_*ρ*_, 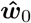). The first three components correspond to each of the three cellular process we aim to model, while the fourth defines the initial conditions. We use Stan’s default Dirichlet prior for the initial condition simplex 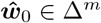. We describe the parameterizations of the remaining three components in the following.

### Cell division

The cell division proportions *δ_i_*(*t*) are parameterized as

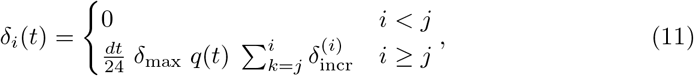

where *δ*_max_ ∈ [0, 24*dt*^−1^] is the maximum division quotient, *q*(*t*) is a function that induces time-dependence in division, and ***δ***_incr_ ∈ Δ^*m*−*j*+1^ is a simplex that defines the relative increase in the division quotient for each size class. For models with time-invariant division (*m*_·*m*·_), *q*(*t*) = 1. The parameter *δ*_max_ is normalized by *dt* in units of days to better facilitate comparisons among models that vary in their values of *dt*; hence, 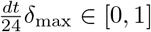. The parameter *δ*_incr_ allows us to constrain cell division to be monotone without imposing a specific functional form of the relationship between cell size and cell division. For models with time-dependent division (*m*_·*t*·_), *q*(*t*) is estimated using a periodic cubic spline with 6 knots and associated control points 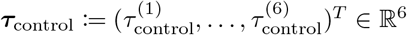. Thus, we have

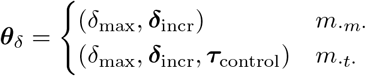

### Carbon fixation

The cell growth proportions are parameterized as

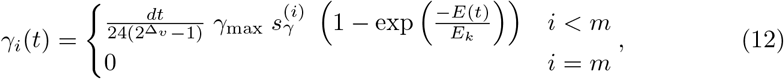

where 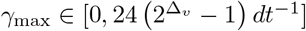 is the maximum cell growth quotient, 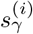 is a function that induces size-dependence in carbon fixation, and *E_k_* ∈ ℝ is a photosynthetic saturation parameter. Recall that *E*(*t*) refers to the incident PAR at time *t*. The parameter *γ*_max_ is normalized by both the choices of time and size discretization to facilitate comparisons between models with different choices of discretization parameters. For models without size-dependent carbon fixation (*m_b_*_··_), 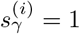. For models with a power-law carbon fixation (*m*_*p*··_),

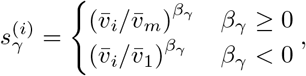

where *β_γ_* ∈ ℝ is a parameter that governs the power-law dependence of carbon fixation on size. For models with a free carbon fixation relationship (*m*_*f* ··_), 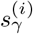 is itself estimated as a parameter separately for each size class. Thus, we have

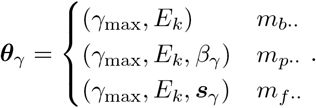

For estimation of the light-saturated photosynthetic rate P_max_, we define the light-saturated growth proportion

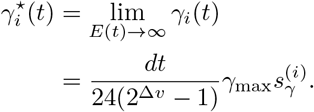

Then, P_max_ is defined as the amount of carbon fixed when *γ_i_*(*t*) is replaced by 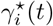 for all size classes *i* and all time points 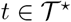.

### Carbon loss

The carbon loss proportions are parameterized as

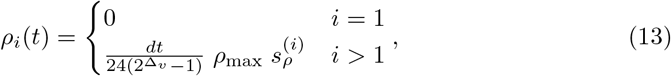

where 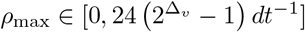 is the maximum cell shrinkage quotient normalized in the same way as γ_max_ and 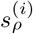 induces size-dependence in carbon loss. For models with no respiration (*m*_··*x*_), 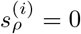. For models with basic respiration (*m*_··*b*_), 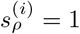. For models with free size-dependent respiration (*m*_··*f*_), 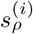 is itself estimated as a parameter as with 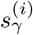. Thus, we have

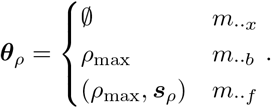

### Experimental data

A publicly available dataset of laboratory experiment time-series measurements of a high-light adapted strain of *Prochlorococcus* [27] collected during the exponential phase of batch growth over two simulated day-night cycles (Fig 2) was used to test model predictions. We used changes in cell abundance over time to calculate division rates, since cell mortality is assumed to be negligible in exponentially growing cultures. A suite of measurements, which include cell size distributions and rates of carbon fixation, were collected at 2 hour intervals for a period of 50 hours to capture two complete diel cycles. Cell size distributions were inferred from flow-cytometry based forward-angle light scatter measurements (FALS). FALS signals normalized by calibration beads were converted to a proxy of mass using the relationships *M* = FALS^1/1.74^ [33] and then converted to carbon quotas assuming an average carbon quotas of 53 fg C cell^−1^ [27]. ^14^C-Photosynthetron experiments were conducted in duplicate at each time point to derive carbon fixation rates, maximum photosynthesis rates, and the photosynthetic saturation parameter. Short (1 hour) incubation times were used to approximate gross carbon fixation rates. Using the 2-hourly cell abundance measurements (*a_t_*), average cell size measurements (*s_t_*) and approximate carbon fixation rates (*f_t_*), we then estimated carbon loss rates (*l_t_*) every 2 hours, using

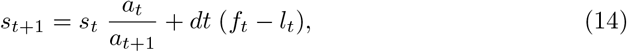

where *dt* is the two hour time step between measurements.

### Implementation

Parameter inference was carried out in the software package Stan [25]. This software performs Bayesian inference, where the target is the *posterior* distribution of the parameters, which reflects the likely values of these parameters given the model, our prior beliefs, and the data [34]. In order to generate samples from the posterior distribution, Stan implements a variant of the Hamiltonian Monte Carlo (HMC) algorithm [35, 36] which has been shown to have superior speed and performance for fitting complex, high-dimensional population dynamics models relative to other Markov Chain Monte Carlo (MCMC) methods for sampling from the posterior [37]. In particular, we use Stan’s implementation of the No-U-Turn Sampler (NUTS) [38] to avoid manual selection of application-specific tuning parameters. Though faster, Stan’s implementation of variational inference provided high instability in model estimates, which may indicate that the approximation to the posterior was of poor quality. Thus, we used HMC, which generated reproducible results and provides asymptotic consistency [36]. The implementation of HMC in Stan uses automatic differentiation to provide the gradients needed to integrate Hamiltonian dynamics. The reader is directed to [39] for additional details on HMC in Stan.

Six HMC chains were run for 2000 MCMC iterations for each model. The 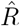 convergence diagnostic [40] was monitored for all model fits to ensure 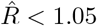, otherwise the sampling procedure was considered divergent.

### Prior distributions

The prior distributions are shown in Table 2. Maximum cell division, carbon fixation and loss along with photosynthetic parameter values were chosen within biologically feasible ranges using information derived from literature [27, 41], otherwise the Stan default priors were used, corresponding to uniform priors [25].

## Acknowledgments

We would like to thank Zachary Johnson for sharing the data. This work was supported by grants from the Simons Foundation (no. 549945 to E.V.A, no. 574495 to F.R., no. 549894 to J.C.) and the Institute for Foundations of Data Science (IFDS; grant no. TRIPODS DMS 2023166 to Z.H.). G.L.B was supported by the Simons Foundation Postdoctoral Fellowship in Marine Microbial Ecology. We also thank Jacob Bien, Chris Edwards and Mick Follows for their support of S.H., J.P.M, and Z.W., respectively, funded by the Simons Foundation (no. 549949 to J.B., C.E. and M.F.). This work was initiated at the Simons Foundation Collaboration on Computational Biogeochemical Modeling of Marine Ecosystems cbiomes.org workshop on Bayesian analysis in marine ecosystems. We thank Helen Hill for workshop organization.

## Supporting Information

### S1 Sliding window experiments

As part of our observation sensitivity experiments, we modified the start time of the model fitting to examine the resulting changes in parameter estimates. As the original cell size distribution dataset only contains two days of data, we appended the dataset to itself to create a 96-hour time series. This allowed us to fit models to a sequence of two-day continuous cell size distribution data that start at different times of the laboratory-simulated light-dark cycle (Fig S1. The start times of these windows ranged from 2 to 46 hours and were spaced four hours apart. In each experiment, the model initialization time is set to match the start time of the window and data outside of the window is discarded.

**Fig S1.**
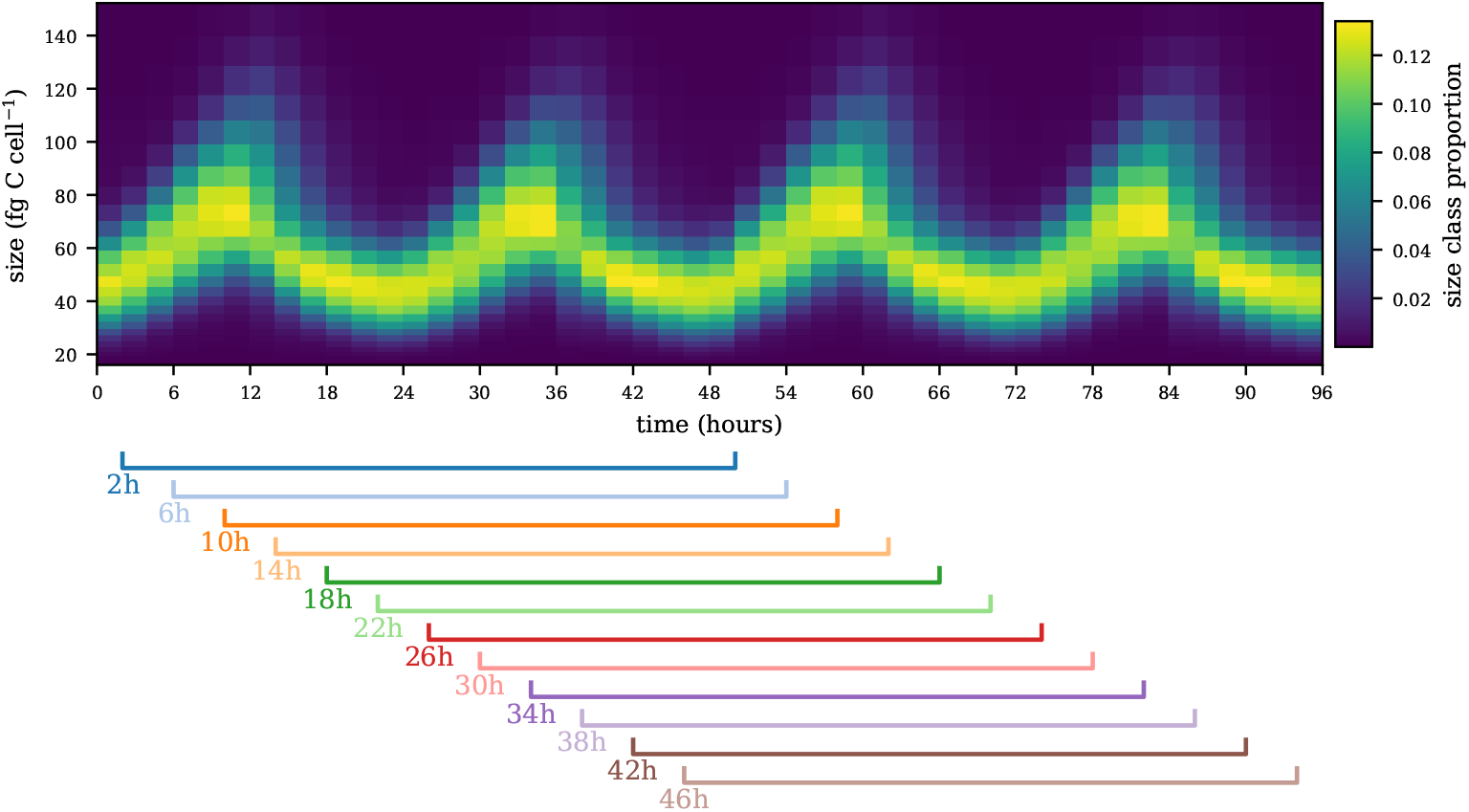
The extended size distribution dataset used in the sliding window experiments and the 2-day windows in which the models are fit.

Here we focus on results for model *m*_bmb_, which are representative for most of our models; when individual models deviate from these results, we note it in the text. Results for all models can be found in the accompanying GitHub repository [1]. Overall, parameter estimates remain consistent for most start times, but we noted a weak cyclical pattern in estimated values and an outlier estimate for a start time of 26 hours (Fig S2), which are both examined below.

The pattern is aligned with the daily cycle and is characterized by increased division and decreased carbon loss rates at start times near 10 hours and – 24 hours later – near 34 hours (Fig S2). It is driven by the estimation of initial conditions at a start time with a large model-observation misfit, which is aligned with the peak of the cell size distribution in most of our models. At the peak of the cell size distribution, for example at t=34 h, the *m*_bmb_ daily cycle underestimates the *Prochlorococcus* cell size distribution (Fig S5). When the estimation window starts at a peak, the estimated initial conditions deviate strongly from the daily cycle steady state solution (compare the solutions of the 2 hours and 10 hours start time at t=10 h, or the 10 hours start time solution at t=10 h and t=34 h; Fig S5D). Due to the increase in the initial cell size distribution, the division rate, which increases with cell size, becomes inflated, impacting other parameter estimates accordingly. This effect is more pronounced for models with a larger model-data discrepancy, while models that fit the size distribution better throughout the daily cycle, such as *m*_ftf_, show a weaker cyclical pattern in the parameter estimates (Fig S3).

**Fig S2.**
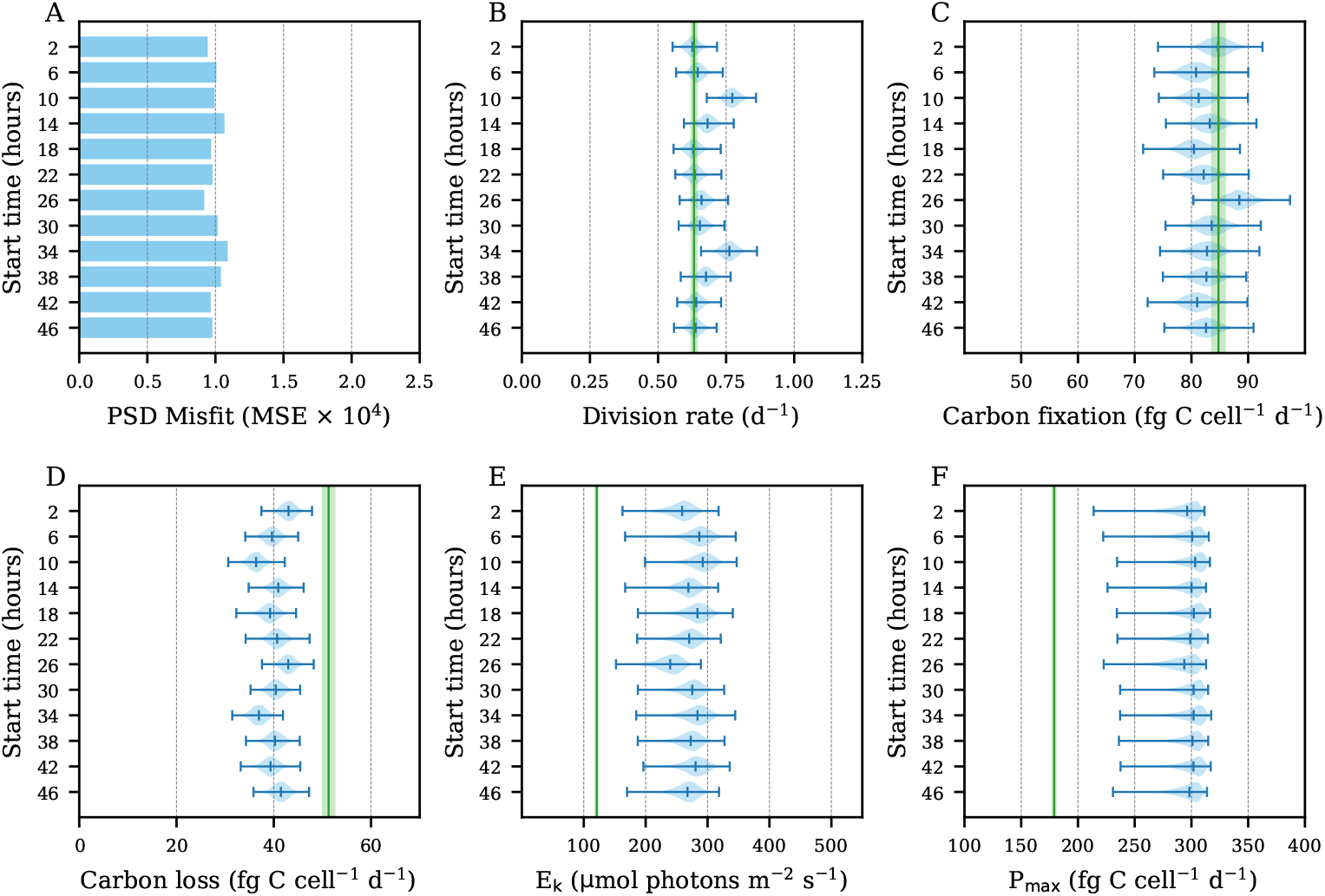
Rate parameter estimates of the model *m*_bmb_ for each window of the sliding window experiment.

Some models, such as *m*_bmx_ and *m*_btb_ (Fig S4), showed much more volatility in their parameter estimates among windows. This indicates that these models may be more unstable and hence their results may be less reliable than the other models.

The second noteworthy pattern in the *m*_bmb_ estimates is the parameter estimate for the start time of 26 hours. Here, the model fitting procedure converged to solutions with higher average carbon fixation and higher carbon loss compared to simulations at other start times. A likely cause for this pattern is the strong correlation structure between the model parameters (Fig 6 in the main document) combined with the broad priors in our model specification. As a result, changes in the start time and associated changes in the order of the observations, in combination with different initial conditions can lead to changes in the posterior estimates that may appear as outliers with respect to the other sliding window experiments. We observed this type of outlier infrequently for most models but it occurred more often for *m*_btb_ which also showed worse convergence properties in our other experiments.

To summarize the stability of our models, we plotted the daily division rate for each model in each window against the concentration parameter *σ* (Fig S6). The vertical spread of each cluster corresponds to the variability of the daily division rate, whereas the horizontal spread corresponds to the variability of the concentration parameter. In general, models with greater values of *σ* exhibited less variability in their daily division rates across windows.

**Fig S3.**
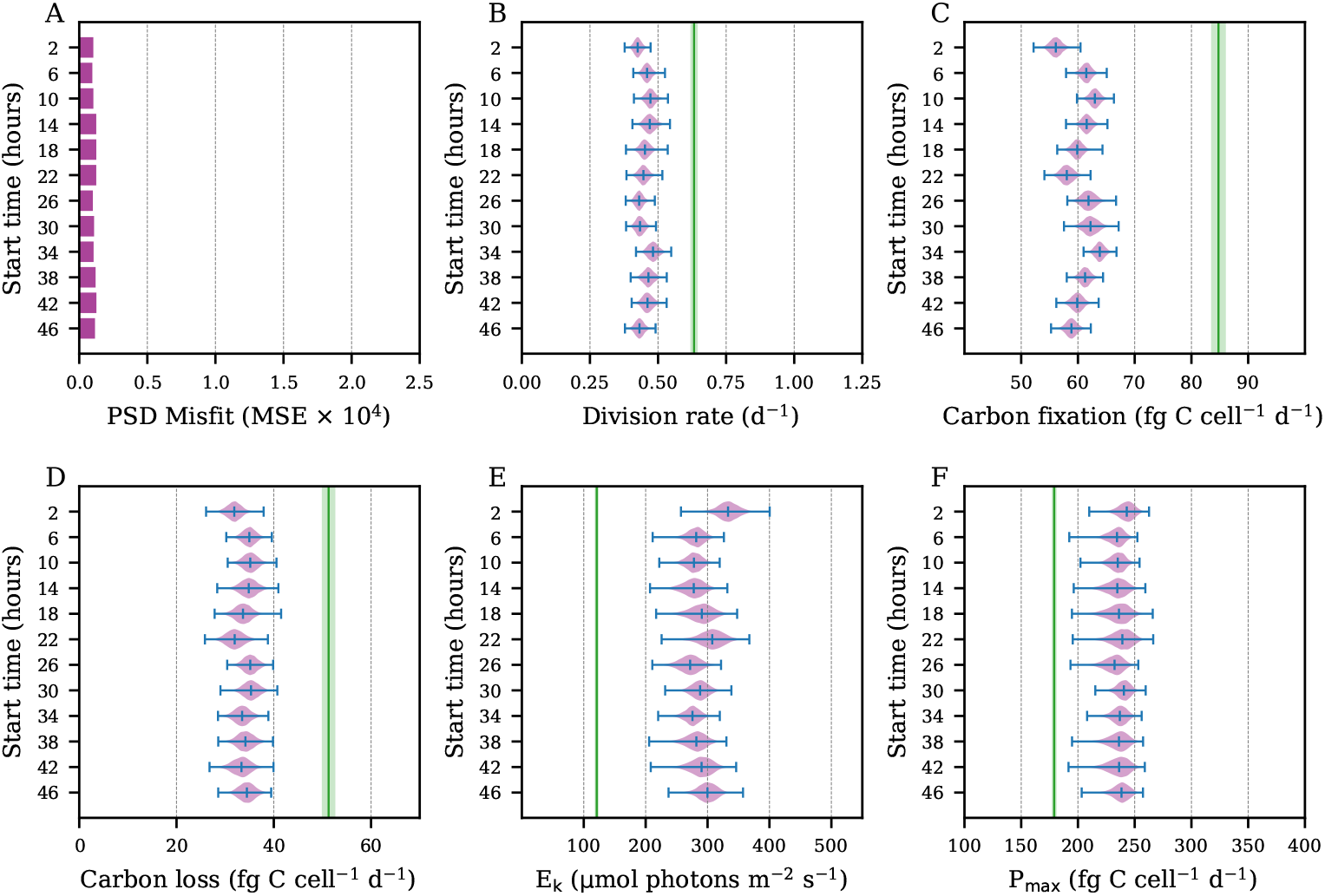
Daily rate parameter estimates of the model *m*_ftf_ for each window of the sliding window experiment. This model showed greater stability in its parameter estimates across windows compared to simpler models such as *m*_bmb_.

**Fig S4.**
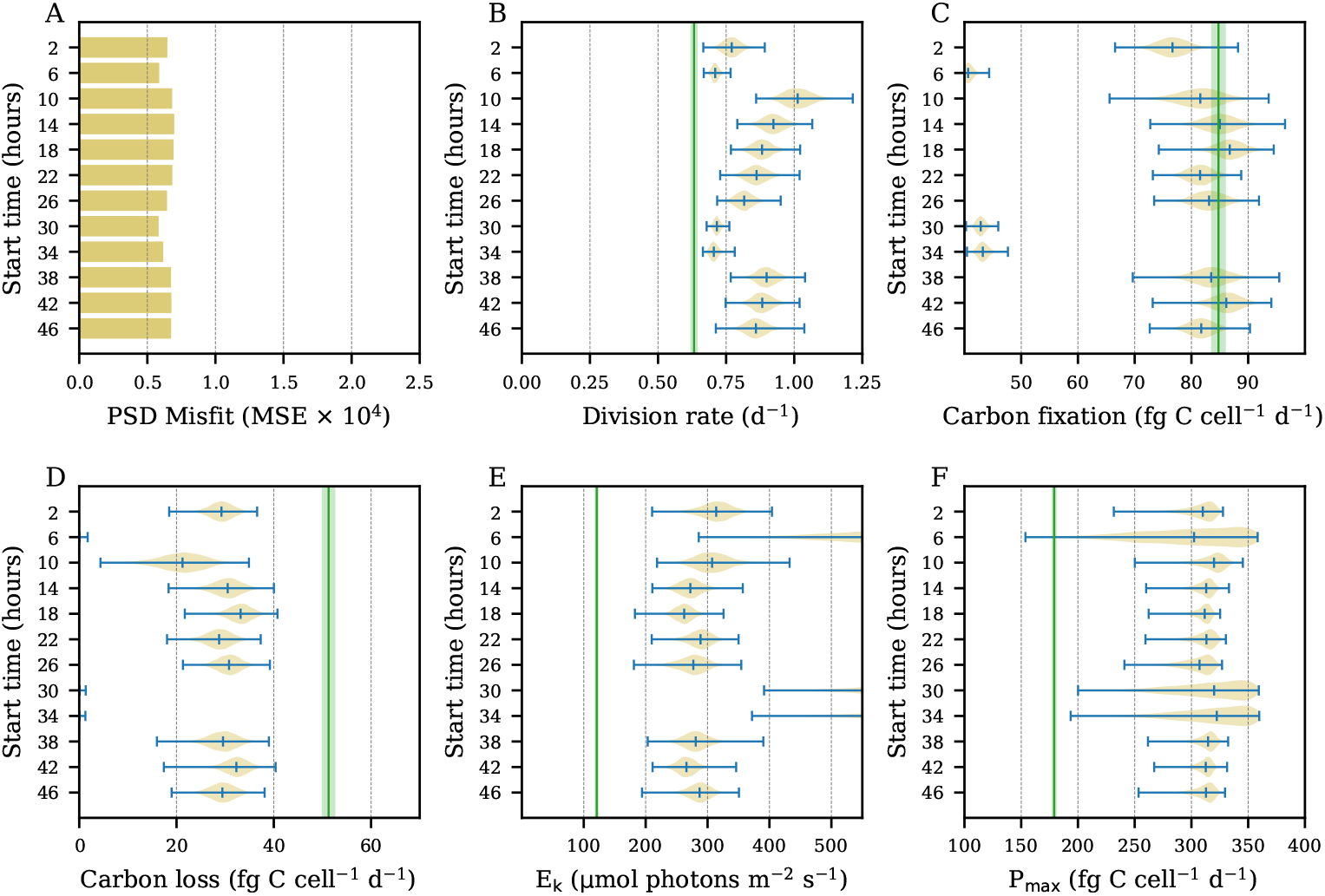
Daily rate parameter estimates of the model *m*_ftf_ for each window of the sliding window experiment. Model results were much more volatile for this model and *m*_bmx_ than the others.

**Fig S5.**
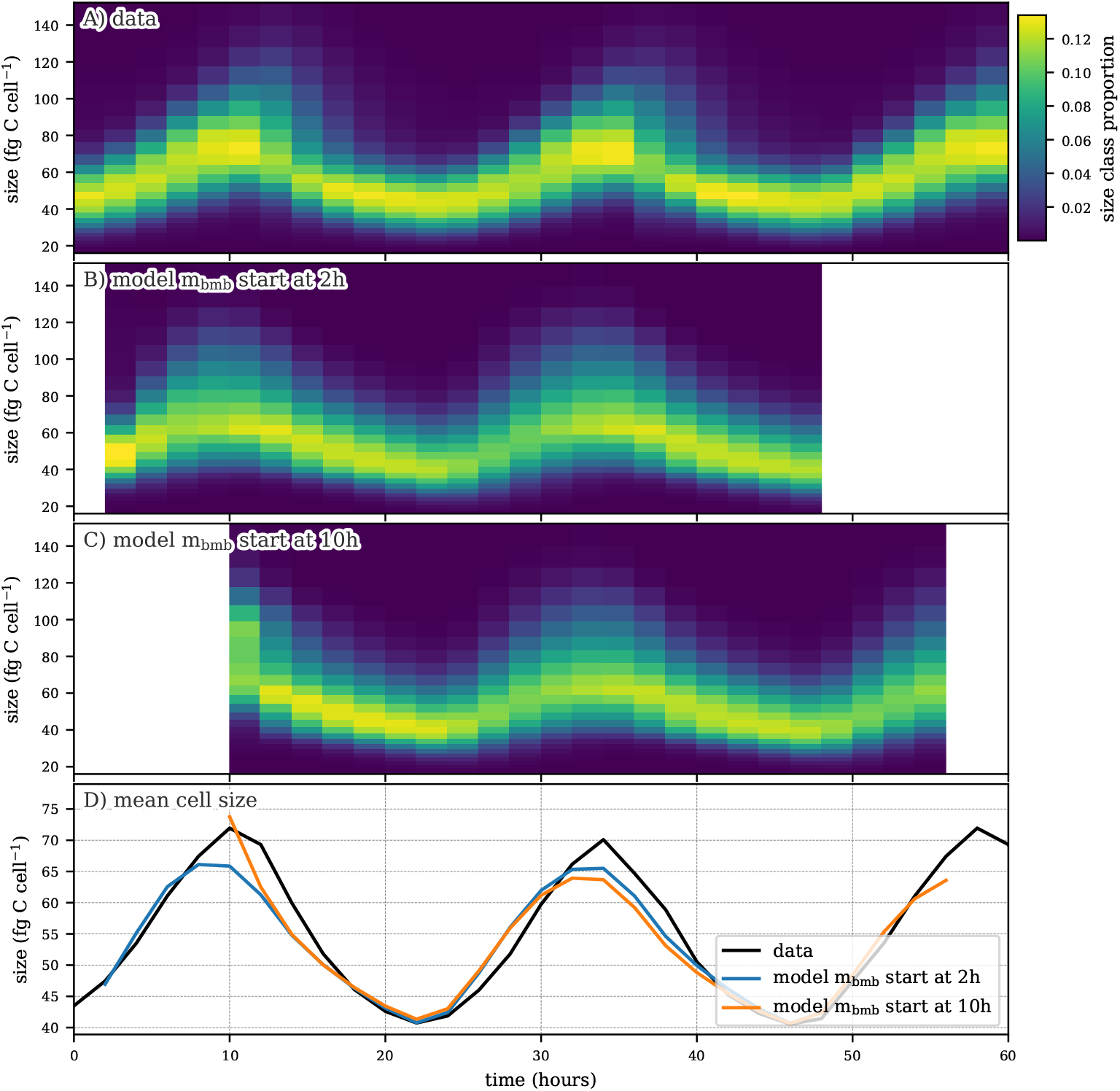
Size distribution in the (A) data, the model *m*_bmb_ in the rolling window experiment started at (B) hour 2 and (C) hour 10. (D) The evolution of the mean cell size in data and model.

**Fig S6.**
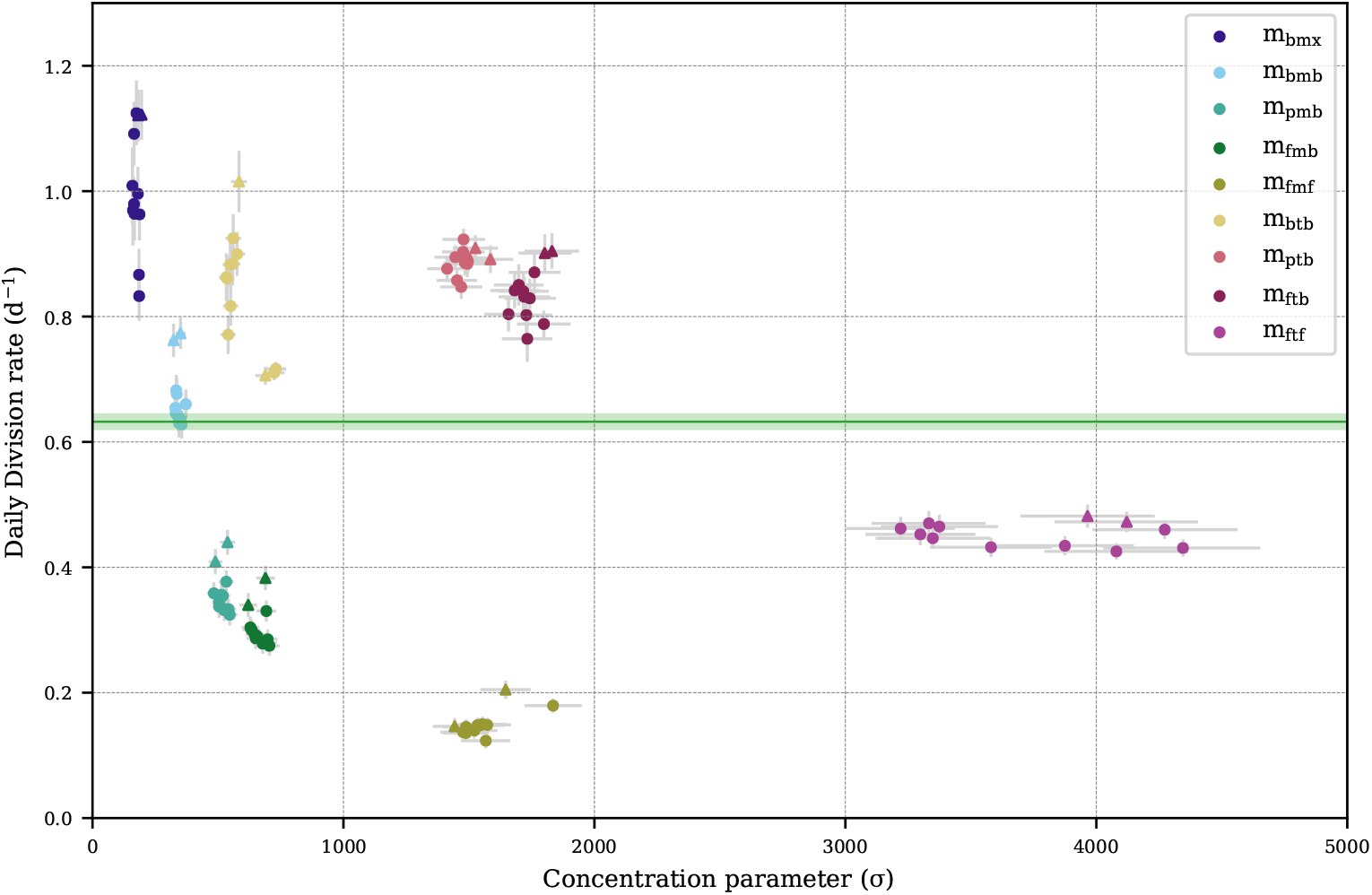
Sliding window experiment daily division rates vs. concentration parameter *σ* by model. Gray error bars indicate one standard deviation of the posterior distribution. Each data point represents the posterior mean daily division rate from one window. Windows starting at *t* = 10 and *t* = 34 are represented as triangles. Green horizontal line indicates observed daily division rate. Green shaded area indicates one standard deviation of uncertainty around the observed value.

**Fig S7.**
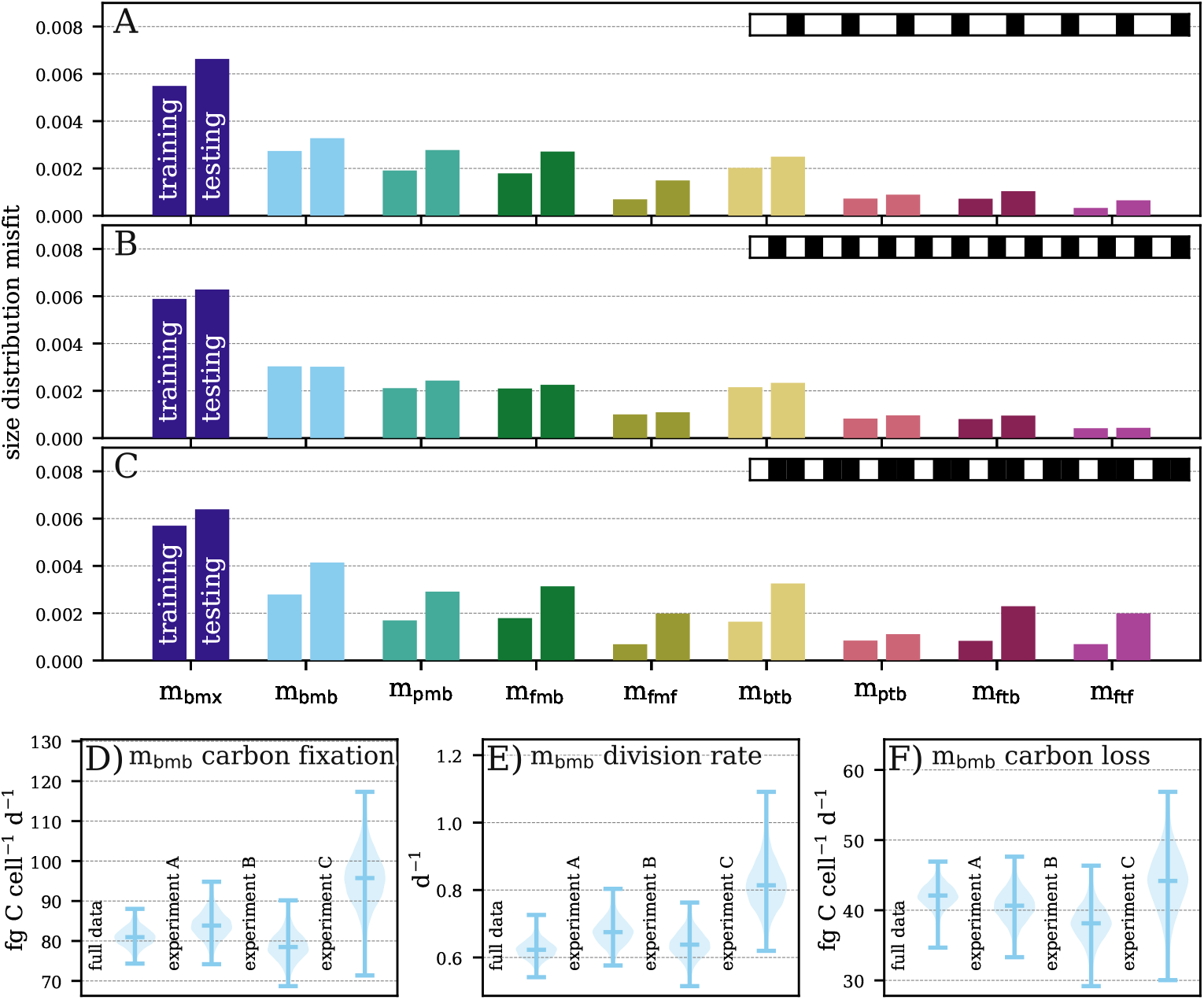
Holdout validation experiments. Size distribution misfit for testing and training data (left and right bar) for each model in the cross-validation experiments (A), (B), and (C) with top right corner visualizing the indices of the testing (black) and training data (white). Examples of the posterior distributions of select model parameters for the full dataset and the two cross-validation experiments: (D) daily carbon fixation rate for *m*_bmb_, (E) daily division rate for *m*_bmb_, and (F) daily carbon loss rate for *m*_bmb_.

### S2 Hold-out validation

In experiment A, the data from every third time step were removed, in experiment B data were removed from every other time step, and in experiment C, two-thirds of the data were removed (see top right corner of Fig S7 A,B,C). As expected, the error on the training data reflected model complexity and decreased from *m*_bmx_ to *m*_fmf_, and again for the models with time-dependent division *m*_btb_ to *m*_ftf_, in all three experiments (Fig S7 A,B,C). While the ratio of testing to training data error increased for more complex models, the absolute value of the testing data error did not increase with model complexity in most of our experiments. The exception involved *m*_ptb_ and *m*_ftb_, which differ only in their size-dependent growth parameterizations. While the more complex *m*_ftb_ with the free growth parameterization exhibited a lower training data error, *m*_ptb_ model with power-law growth achieved a lower testing data error. Taken together with the results for *m*_pmb_, which were similar to those of *m*_fmb_, we have some evidence that the power-law growth parameterization is suitable for models in this application, creating a size-dependent growth relationship that performed better on testing data than a freely estimated relationship.

Reducing the number of observations in the training set had a noticeable impact on the models parameter estimates (Fig S7 D-F). With less data in the training dataset, the posterior distributions of the estimated parameters broadened from those obtained using the full dataset and eventually showed shifts in the mean parameter estimates when more data is excluded (e.g. *m*_bmb_ daily division in experiment C, Fig S7 E). The broadening matches our intuition: fewer observations constrain the parameter estimates to a lesser extent than the information contained in the full dataset. With two thirds of the data excluded and observations occurring every 6 hours, the rate parameters could no longer be estimated reliably and mean parameter estimates deviated noticeably from their values on the full dataset. In summary, when as much as one half of the data was removed, the predicted rate parameters still capture the daily cycle of *Prochlorococcus* dynamics. Estimates for the parameters of interest also remained stable.

### S3 Hourly rate estimates

Here, we show the hourly rate estimates for all nine models (Fig S8). The trends discussed in the main text can be seen in the 4 remaining models (*m*_pmb_, *m*_fmb_, *m*_btb_, *m*_ftb_). Models *m*_pmb_ and *m*_fmb_, which assume cell division only varies as a function of cell size, predicted cell division to occur too early (Fig S8 A). Again, model *m*_ftb_, with both time-dependent division and size-dependent carbon fixation, correctly predicted the timing of cell division, but overestimated division during the morning. The model with time-dependent division but no size-dependence in carbon fixation (*m*_btb_) did not correctly predict the timing of cell division. All models underestimated carbon fixation (Fig S8 B) as seen in the main text. They also overestimated carbon loss at night and underestimated carbon loss during the day (Fig S8 C).

Of the four models we exclude from the figure in the main text, models that overestimated cell division rates (*m*_btb_, *m*_ftb_) predicted higher proportions of dividing cells for smaller sizes (Fig S8 D). Similarly, the models that underestimated cell division (*m*_pmb_, *m*_fmb_) predicted very low proportions of dividing cells in the large size classes. As with the other models that assume no size dependence in carbon fixation, *m*_btb_ estimated the maximum possible fraction of dividing cells to be near 1 (Fig S8 E). Again, models with a size-dependent carbon fixation parameterization (*m*_pmb_, *m*_fmb_, *m*_ftb_) predicted the maximum proportion of growing cells to decrease as the size of the cells increased. The predicted fractions of cell shrinkage tended to be significantly lower than the fractions of maximum growth, ranging from negligible to about one-fifth of the peak maximum growth fraction (Fig S8 E, F), as observed in the main text.

---

**Fig S8.**
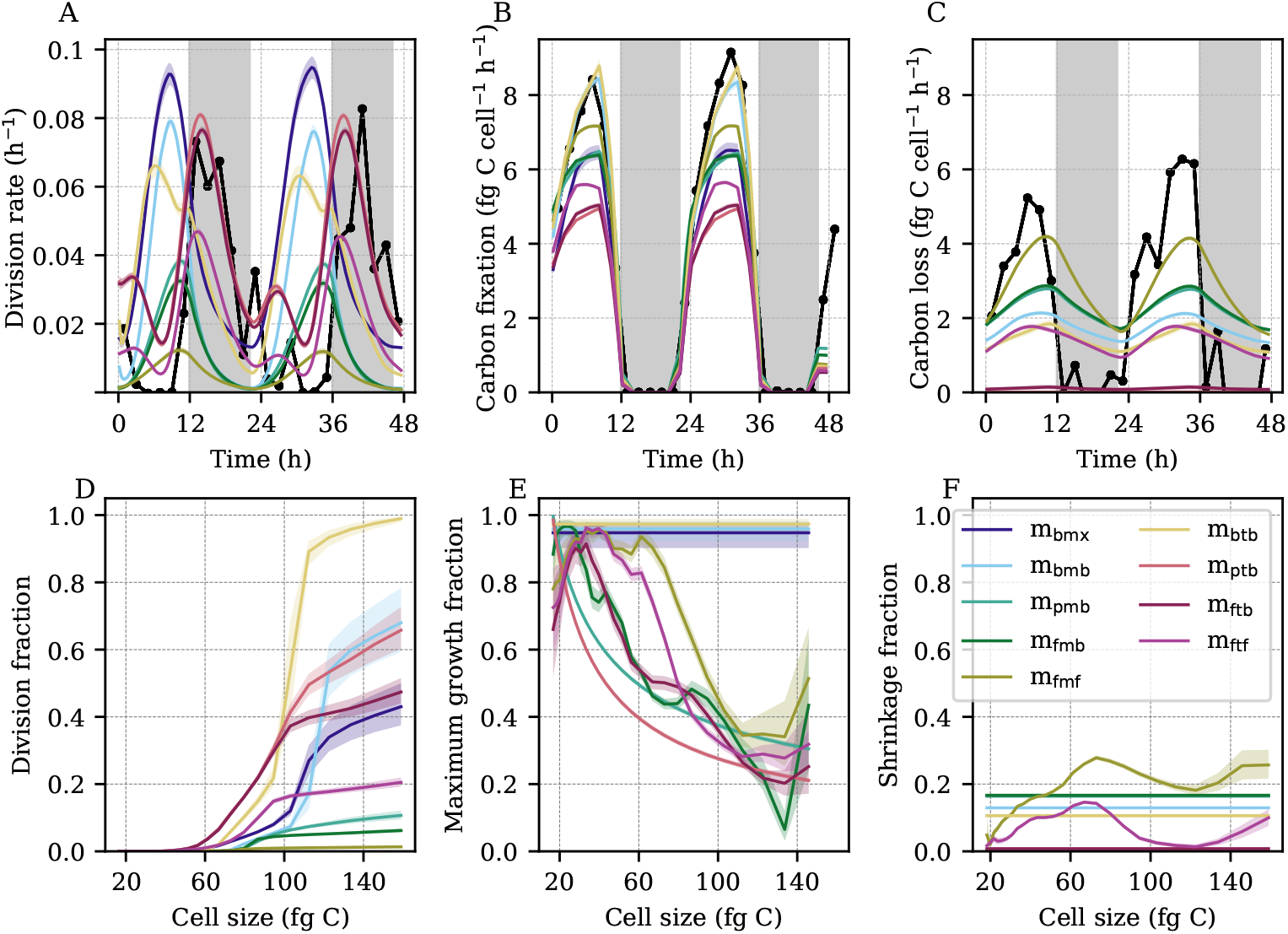
Model predicted hourly rate parameters. (A) Observed (black) and predicted (colored bands) hourly division rates. (B) Observed (black) and predicted (colored bands) hourly carbon fixation. (C) Observed (black) and predicted (colored bands) hourly carbon loss. (A-C) Black points indicate ground truth calculated from data. (D) Predicted cell division fraction as a function of cell size. (E) Predicted light-saturated cell growth (carbon fixation) fraction as a function of cell size. (F) Predicted cell shrinkage (carbon loss) fraction as a function of cell size. (A-F) Colored bands indicate model estimates. Shading indicates the first to third quartiles of the posterior distributions. (D-F) Fractions correspond to MPM transitions over a 20-minute time period.

## Notes

https://github.com/CBIOMES/bayesian-matrix-population-model

### Competing Interest Statement

The authors have declared no competing interest.

